# Involvement of the putative metal efflux protein YbeX in ribosomal metabolism

**DOI:** 10.1101/2023.03.20.533420

**Authors:** İsmail Sarıgül, Amata Žukova, Emel Alparslan, Margus Pihlak, Sille Remm, Tanel Tenson, Ülo Maiväli

**Affiliations:** University of Tartu, Institute of Technology, Nooruse 1, 50411, Tartu, Estonia; Tallinn University of Technology, Department of Cybernetics, Ehitajate tee 5, 19086, Tallinn, Estonia; University of Zurich, Institute of Medical Microbiology, Gloriastrasse 28/30, 8006, Zürich, Switzerland

## Abstract

YbeX of *Escherichia coli*, a member of CorC protein family, is a putative Co^2+^/ Mg^2+^ efflux factor. Here, we describe several *ΔybeX* phenotypes and report an involvement of YbeX in ribosomal metabolism. *E. coli* lacking *ybeX* has a longer lag phase on outgrowth from the stationary phase. This phenotype is heterogeneous at the individual cell level and can be rescued by supplementing the growth media with magnesium. *ΔybeX* strain is sensitive to elevated growth temperatures and to several ribosome-targeting antibiotics, which have a common ability to induce the cold shock response in *E. coli*. *ΔybeX* cells accumulate distinct 16S rRNA degradation intermediates present in both 30S particles and 70S ribosomes. We propose that a function of YbeX is maintaining the magnesium homeostasis in the cell, which is needed for proper ribosomal assembly.

## INTRODUCTION

Ribosomal biogenesis is a highly regulated process which encompasses concomitant transcription, processing, degradation, modification and folding of ribosomal RNAs, as well as equimolar synthesis and incorporation into ribosomes of more than 50 different ribosomal proteins (Davis and Williamson, 2017). In bacteria, this is catalyzed, chaperoned and generally facilitated by dozens of dedicated proteins working in tandem in several partially overlapping and redundant pathways (Shajani *et al*., 2011). However, due to its sheer complexity, our understanding of this process is bounded to isolated fragments of processing/folding pathways, with minimal knowledge of many individual factors’ precise mechanisms of action.

It has long been known that Mg^2+^ is necessary for ribosomal assembly and translation (McCarthy *et al*., 1962). More recently, it was discovered that intracellular free Mg^2+^ and rRNA transcription are actively co-regulated for achieving optimal ribosomal assembly and translation (Pontes *et al*., 2016). Also, it has been shown that Mg^2+^ influx can provide an active mechanism to alleviate ribosomal stress phenotypes, probably by stabilizing ribosomal structure (Lee *et al*., 2019).

*ybeX* encodes a putative Co^2+^/Mg^2+^ efflux protein, which is highly conserved in bacteria, but poorly characterized (Kazakov *et al*., 2003; Anantharaman and Aravind, 2003). In the genome of *E. coli*, it is located in the *ybeZYX-Int* operon (**Fig. 1A**), transcripts of which have not been fully mapped. The *lnt* gene, which encodes an essential inner membrane protein, is predicted to be under the control of the minor heat shock sigma factor σ^24^ (RpoE) (Keseler *et al*., 2013), while transcription of *ybeY*, *ybeZ* and *ybeX* is regulated by the primary heat shock sigma factor σ^32^ (RpoH) (Nonaka *et al*., 2006). In low-magnesium conditions, the levels of YbeX (but not of YbeY and YbeZ) mRNA and protein are about two-fold reduced, consistently with its proposed role in Mg^+2^ efflux (Caglar *et al*., 2017).

**Fig. 1.**
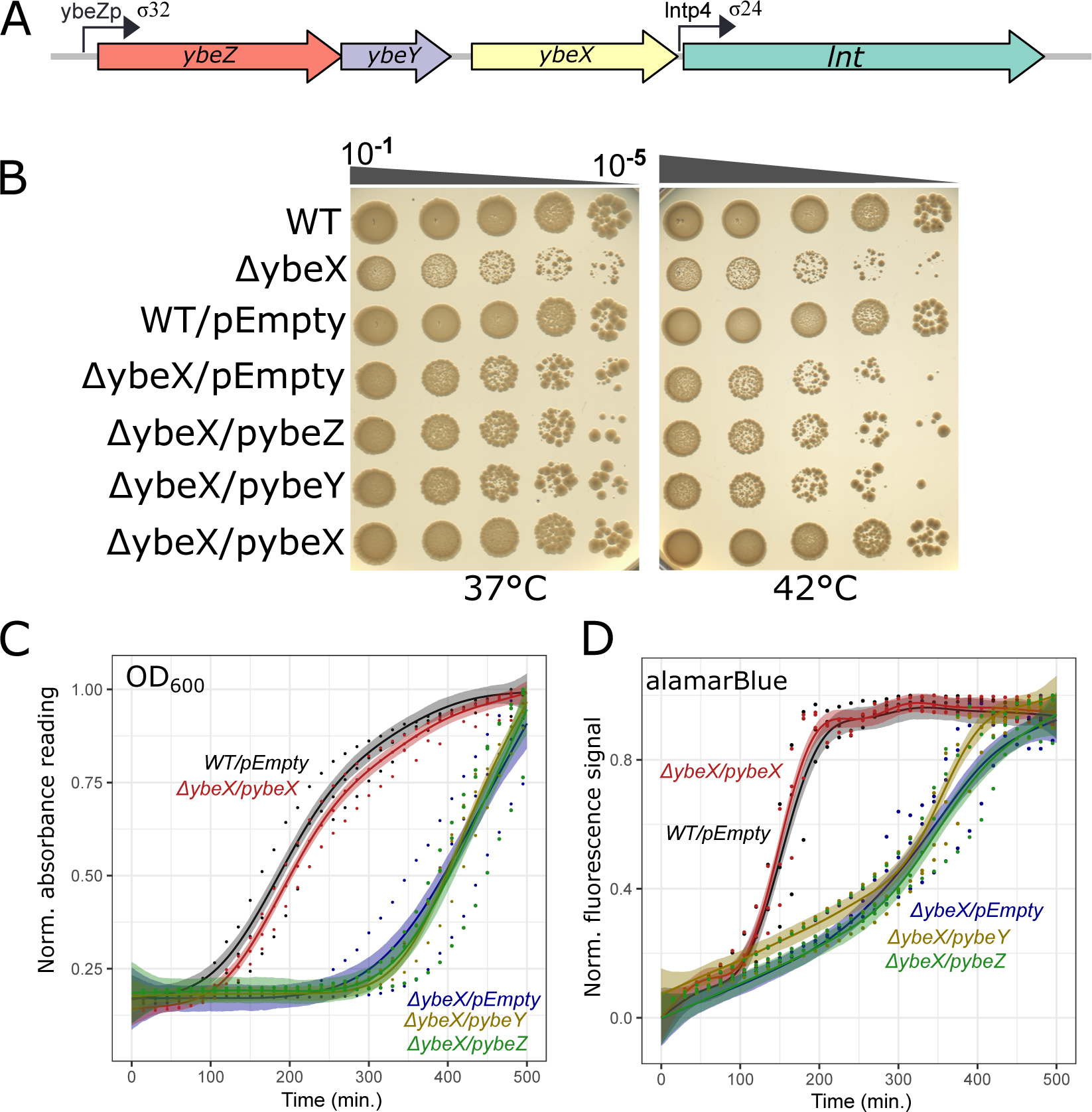
Growth phenotypes of *ΔybeX* strain and compensation with single-copy plasmid. (**A**) The *E. coli ybeZYX-lnt* operon chromosomal organization with designed sigma factors (σ^32^ and σ^24^). (**B**) Dot spot assay with wild type (WT) and *ΔybeX* strain, along with the wild type strain harboring empty plasmid (WT/pEmpty), *ΔybeX* strain transformed with empty plasmid, and *ΔybeX* strain conjugated with YbeZ, YbeY, or YbeX-expressing single copy TransBac library plasmid. LB agar plates without any antibiotics were incubated at 37°C or 42°C. (**C-D**) The stationary phase outgrowth of wild type and *ΔybeX* strains harboring empty plasmid in liquid LB medium. Panel C shows the OD600 signal, and panel D shows the alamarBlue fluorescence reading normalized to one. Individual measurements from independent experiments, each presented as the mean value of three technical replicates, are shown as dots. Curves are presented as modelled splines, and 95% credible intervals are shown as shaded areas (see Materials and methods for details). pybeX, pybeZ and pybeY denote the YbeX, YbeZ and YbeY expressing plasmid.

The most-studied member of the *ybeZYX-lnt* operon is the *ybeY*, whose importance in ribosomal metabolism is beyond dispute, while the precise mode of action of YbeY remains unclear (Davies *et al*., 2010). The YbeY is, by sequence homology and structural studies, a zinc-dependent RNA endonuclease. YbeY is universally conserved over the three domains of life, has very strong, albeit heterogeneous, phenotypes in every organism that has been looked into, and it has been shown by genetical methods to be required for the correct processing of the 3’ end of 16S rRNA (Liao *et al*., 2021). Moreover, YbeY mutants have been shown to be defective in translation and accumulate defective ribosomes in several bacterial species, mitochondria and chloroplasts (Summer *et al*., 2020; Liao *et al*., 2021; D’Souza *et al*., 2021). And yet, in the purified form, its RNase activity seems to be limited to short RNA oligonucleotides (Jacob *et al*., 2013; Babu *et al*., 2020), while *in vitro* processing of the 16S rRNA 3’-end can be achieved without it (Smith *et al*., 2018).

The *ybeZ* gene is located upstream of *ybeY*, having four nucleotides overlap. *ybeZ* encodes a phosphate starvation-regulated PhoH subfamily protein with the NTP hydrolase domain (Kim *et al*., 1993). YbeZ has phosphatase activity and is a putative RNA helicase through sequence homology (Kazakov *et al*., 2003; Andrews and Patrick, 2022). A physical interaction between YbeY and YbeZ was suggested based on bacterial two-hybrid system experiments in *E. coli* (Vercruysse *et al*., 2016). Their interaction has been biochemically verified in *Pseudomonas aeruginosa* (Xia *et al*., 2020).

Here we characterize the growth and ribosomal homeostasis phenotypes of the deletion of *ybeX* in *Escherichia coli*.

## RESULTS

### Deletion of *E. coli ybeX* leads to heat sensitivity and longer outgrowth from the stationary phase

*ybeX* is a part of the RpoH (heat response) regulon (Nonaka *et al*., 2006). We tested by a spot assay the effect of elevated growth temperature on the *ΔybeX* strain from the Keio collection (Baba *et al*., 2006), compared to the isogenic BW25113. After overnight growth in the LB liquid medium, serial dilutions of the culture were spotted on LB agar plates and incubated at 20°C, 37°C or 42°C overnight. Disruption of *ybeX* hindered growth at 42°C but not at 20°C (**Fig. 1B**, **Fig. S1a**). For verification, *ybeX* deletion was reintroduced in two strain backgrounds, MG1655 and BW25113. We verified the deletion of *ybeX* and the presence of kanamycin resistance cassette by PCR analysis (**Fig. S1b,c**). Heat sensitivity occurred in both newly constructed *ΔybeX* strains (**Fig. S1d).** This demonstrates that the observed phenotype is *ybeX*-inflicted. We used the *ybeX* deletion strain of the Keio collection in further studies.

Next, we assessed whether the lack of the YbeX protein caused heat sensitivity. Alternatively, secondary effects of the chromosomal deletion might be responsible for this phenotype. We reintroduced *ybeX* on a single-copy TranBac library plasmid (Otsuka *et al*., 2015) and found the leaky YbeX expression in the absence of the inducer (isopropyl-β-D-1-thiogalactopyranoside; IPTG) was sufficient to rescue the heat sensitivity of the *ΔybeX* mutant.

The empty vector (pEmpty) and TranBac plasmids carrying *ybeY* or *ybeZ* had no effect on the growth (**Fig. 1B**). Thus, the heat sensitivity of the *ΔybeX* strain was caused by the absence of the YbeX protein, rather than through polar effects on neighboring genes.

To find which growth phase is affected by the *ybeX* deletion, we monitored bacterial cultures in liquid LB medium at 37°C on a 96-well plate reader. We did not notice differences between the growth of WT and *ΔybeX* strains when cultures were started from freshly grown single colonies (data not shown). When cultures were inoculated with bacteria from the stationary phase overnight cultures, the *ΔybeX* mutant had a much longer lag phase (300-350 min.) compared to the WT (100-150 min.; **Fig. 1C**, **Fig. S2a**). Both strains reached the same optical density in the stationary phase. A similar number of colonies after dilution and plating WT and *ΔybeX* (**Fig. 1B**) indicates that the delay of the visible growth of the *ΔybeX* mutant is not caused by decreased survival in the stationary phase but reflects later regrowth of the same number of live bacteria. Expression of *ybeX* from a single-copy plasmid abolished the prolonged lag phase completely while complementation with the plasmids carrying either *ybeY*, *ybeZ* or *lnt* had no effect confirming that lack of the YbeX protein is causing the delay of regrowth, while further excluding polar effect as a cause of the *ΔybeX* phenotype (**Fig. 1C; Fig. S2b**).

To investigate whether the longer lag phase of *ΔybeX* strain is due to lower metabolic activity in the mutant cells, we used the alamarBlue reagent, a quantitative indicator of the oxidation-reduction potential of cell membranes, as a proxy for metabolic activity (Rampersad, 2012). In a negative control experiment conducted in PBS buffer lacking the nutrients necessary for the resumption of growth, both strains show similarly low alamarBlue signal, indicating similar levels of metabolic activity (the superimposed black lines in **Fig. S2c**). When diluted into fresh LB medium, the alamarBlue signal immediately starts to increase for both strains, indicating activation of similar levels of cellular metabolism (**Fig. S2d,e**). While the initial rate of increase in the alamarBlue signal, and by implication the cellular metabolism levels, are equal for WT and *ΔybeX* cells, after about 100 minutes the WT acquires a still faster rate of signal increase, while the *ΔybeX* cells continue as before for about 200 more minutes, before the rate of their signal growth increases to WT levels (**Fig. S2d**). As shown by the OD_600_ measurements (**Fig. S2a**), for both the WT and the *ΔybeX* cells, this phase shift in redox power is accompanied by the start of cell divisions (**Fig. S2e**). These results indicate that the longer lag phase of the *ΔybeX* strain is not caused by lower levels of metabolic activity in the *ΔybeX* cells whilst they are preparing for the resumption of cell divisions. Nor is it caused by a later onset of said metabolic activity.

### The delayed outgrowth of *ΔybeX* is heterogeneous at the individual cell level

When streaking out mutant strains from glycerol stocks and overnight grown stationary phase cultures, we noticed that while the *ΔybeZ* strain produces wild-type-like colonies and *ΔybeY* strain has uniformly small colonies, the *ΔybeX* strain produces heterogeneous colonies. Re-streaking of small and large *ΔybeX* colonies resulted in well-grown large second-generation colonies, indicating that the heterogeneous phenotype is not caused by a genetic mutation (data not shown). We also tested whether the colony heterogeneity is caused by freeze-thawing in the glycerol mix by growing the *ΔybeX* and wild-type cells in liquid media into stationary phase and then plating the cells directly onto LB agar plates. Again, while wild-type cells exhibited uniform colonies, the *ΔybeX* strain gave heterogeneous colony growth (**Fig. 2A**). Thus, it is likely that the growth heterogeneity of *ΔybeX* depends on the heterogeneity of the initial physiological states of individual stationary cells.

**Fig. 2.**
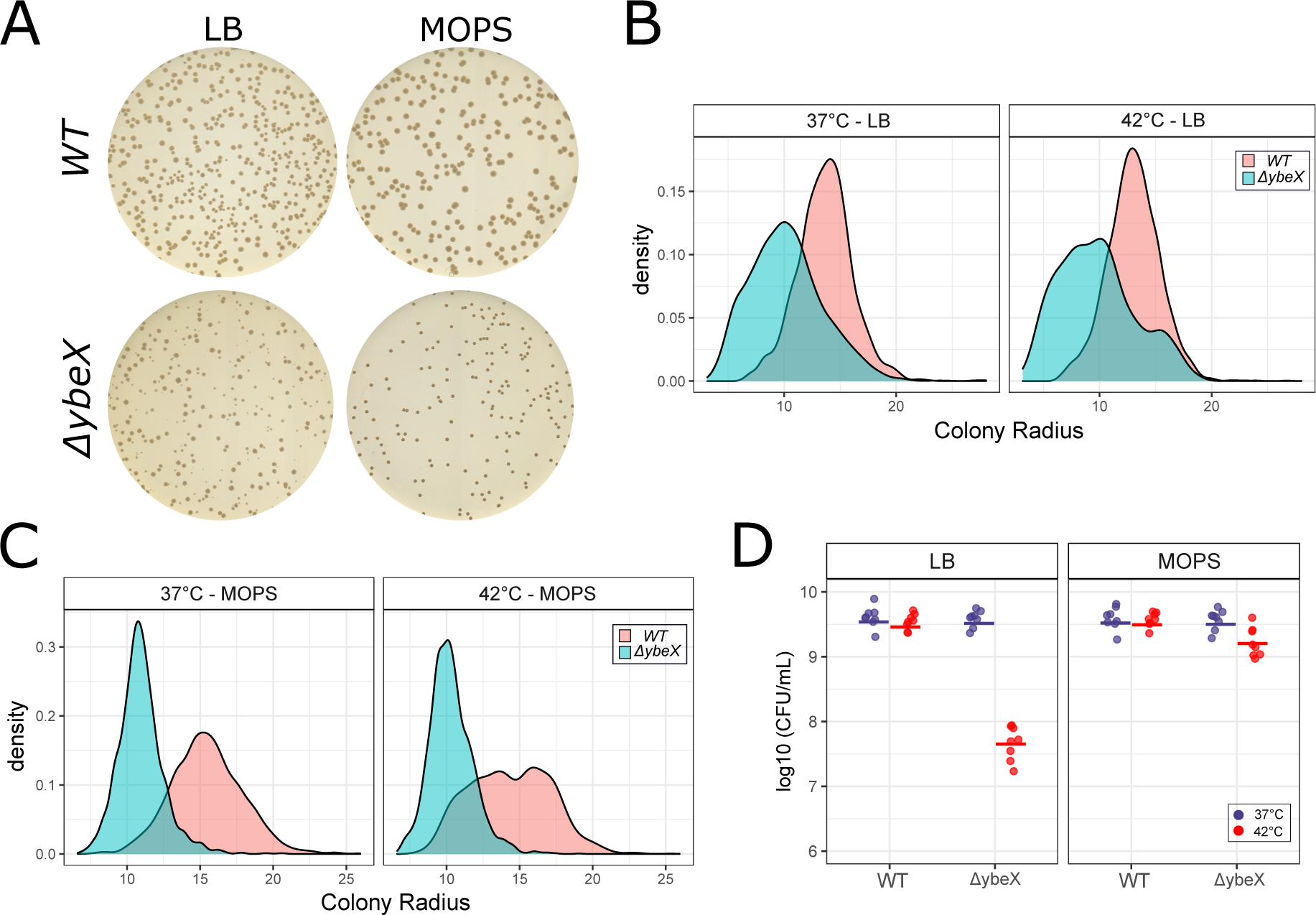
Characterization and quantification of *ΔybeX* and Keio wild-type (WT) strains colony sizes at 37°C and 42°C. (**A**) Visual inspection of colony appearance of WT and ΔybeX strains on LB agar plates. The cells were grown in LB liquid medium or MOPS minimal medium, serially diluted, and plated on LB agar plates. The plates were incubated at 37°C overnight. (**B**) Estimated colony counts for WT and *ΔybeX* strains grown in LB or MOPS MM. Diluted cells were plated on LB Agar plates and incubated overnight at 37°C and 42°C. (**C**) Density plots of the distribution of quantified colony radiuses of *ΔybeX* and isogenic WT strains at 37°C and 42°C.

We quantified colony radiuses of wild-type and *ΔybeX* strains grown in LB and MOPS minimal media supplemented with 0.3% glucose using AutocellSeg (Khan *et al*., 2018). *ΔybeX* cells tend to form smaller colonies than wild-type cells when grown in LB and MOPS media (**Fig. 2A**). *ΔybeX* colonies are heterogeneous when grown in LB medium (**Fig. 2B**), while the colony radiuses are homogeneous when *ΔybeX* cell s are grown in MOPS liquid medium (**Fig. 2C**).

We then asked how heat shock affects cell growth and heterogeneity. Overnight cultures were diluted and plated on LB agar plates following 16-18 hours of incubation at 37°C or 42°C (**Fig. S3a**). We observed fewer *ΔybeX* colonies at 42°C (p < 0.0001 and p = 0.02 for LB and MOPS, respectively; **Fig. 2D**).

We inspected the colony growth of *ΔybeX::kan* (*ybeX* single deletion strain in BW25113 background constructed via lambda red recombination) and *ΔybeX/kan-* (the kanamycin cassette removed from the inhouse constructed *ΔybeX::kan*) cells at 37°C for 24 and 48 hours (**Fig. S3b,c**). No significant differences were observed for *ΔybeX::kan* and *ΔybeX/kan-*. Furthermore, although the observed tiny colonies of *ΔybeX* were increasing in size over time, they consistently remained smaller than WT-like *ΔybeX* colonies (**Fig. S3b,c**).

To better understand the nature of the observed lag phase phenotype at the individual cell level, we quantified colony radiuses of the *ΔybeX* and the WT cells at 37°C and 42°C using cells pre-grown for 16-18 hours in liquid LB or MOPs minimal media prior to plating. Inspection of four independent stationary phase outgrowth experiments showed, in accordance with our previous observations, that at both temperatures WT cells tend to form larger colonies, while the colony radiuses of *ΔybeX* cells are heterogeneous and possibly dimorphic. These intuitions were formalized by jointly modelling means and standard deviations of colony radiuses and, in a separate model, the colony radiuses as mixtures of two normal distributions (see Materials and Methods for details). The estimated mean colony radiuses are smaller in the ybeX strain by about 1/3 (the difference, in arbitrary units, at 37°C is 3.58 [95% CI 2.89, 4.24] and at 42°C is 3.72 [3.01, 4.42]), and the *ΔybeX* colony radiuses have a larger standard deviation (the difference in 37°C is 0.67 [0.36, 1.04] and at 42°C is 0.91 [0.53, 1.34]). Interestingly, modelling the colony radiuses as emanating from two distinct gaussian populations resulted in a superior out-of-sample fit, as assessed by leave-one-out-cross validation (data not shown), supporting the conjecture that *ΔybeX* cells grow in at least two distinct regimes, one of which is similar to WT growth, while another results in up to two-fold smaller colonies (WT vs. *ΔybeX* difference in 37°C: μ_1_ (estimate for the mean of the first gaussian): 1.73 [0.47, 2.93], μ_2_: 5.58 [4.71, 6.39], and in 42°C: μ_1_:1.39 [0.06, 2.81], μ_2_: 5.92 [4.99, 6.79]).

### The *ΔybeX* strain is sensitive to ribosome-targeting antibiotics

*ybeX* disruption has been reported to cause cell death in the presence of chloramphenicol (Smith *et al*., 2007). We therefore explored the effects of various antibiotics on the *ΔybeX* cells. First, we determined the minimal inhibitory concentrations (MICs) in LB for the *WT* and the *ΔybeX* strains (see Materials and Methods). The MICs were two times lower for *ΔybeX* in the presence of fusidic acid, clindamycin, chloramphenicol, tetracycline and erythromycin (**Table S1**). These structurally unrelated ribosome-targeting antibiotics have been shown to induce cold-shock proteins or block the induction of heat-shock proteins (VanBogelen and Neidhardt, 1990; Cruz-Loya *et al*., 2019). We further inspected the effects of these antibiotics using the dot spot assay described above, except that the LB agar plates were supplemented with sub-inhibitory concentrations of indicated antibiotics (see Materials and Methods). The *ΔybeX* strain exhibited severe sensitivity to sub-lethal concentrations of all of these antibiotics (**Fig. 3A**). Expressing the YbeX protein from a single-copy plasmid in the absence of the inducer completely rescued the antibiotic sensitivity (**Fig. 3B**). In contrast, protein synthesis-targeting antibiotics, for which we do not have evidence that they affect the cold shock response (amikacin, streptomycin, kanamycin, tobramycin and mupirocin), was founded not to have an effect. In addition, the transcriptional inhibitor rifampicin revealed no effect on the *ΔybeX* strain compared to the WT (**Fig. 3**). We also measured the MICs in the MHB cation-adjusted media. The MICs for wild type and *ybeZ*, and *ybeX* deletion strains remained the same, while *ybeY* deletion strain showed lower MICs for all tested antibiotics (data not shown).

**Fig. 3.**
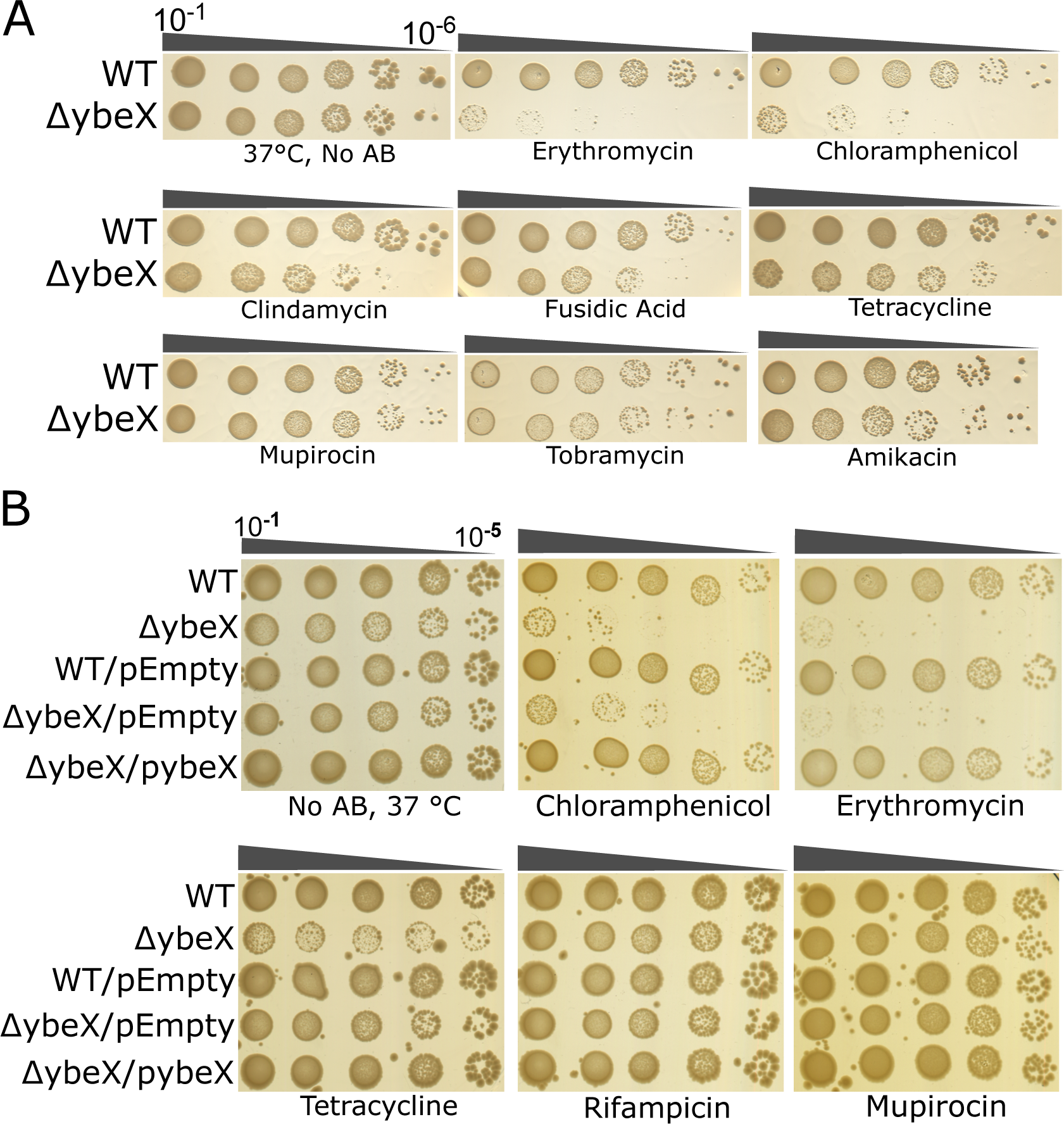
*ΔybeX* cells exhibit severe sensitivity to sub-lethal concentrations of ribosome-binding antibiotics. (**A**) Wild-type BW25113 strain and *ΔybeX* cells were grown overnight in LB liquid medium, serially diluted and spotted on LB agar plates supplemented with sub-inhibitory concentrations of indicated antibiotics or without antibiotic (No AB). The plates were incubated at 37°C overnight. (**B**) Representative plates from a dot spot assay with strains described in Fig. 1 are presented. pybeX, pybeZ and pybeY denotes the YbeX expressing plasmid.

We also tested the survival of two isogenic wild-type strains, MG1655 and BW25113 and the corresponding deletion strains *ΔybeX::kan^MG^* and *ΔybeX::kan^BW^* under sub-inhibitory antibiotic concentrations. Both genetic backgrounds exhibited similar antibiotic sensitivities, and removal of the kanamycin resistance cassette (in strains *ΔybeX/-kan^MG^* and *ΔybeX/-kan^BW^*) had no effect (**Fig. S4**). In contrast, ectopic expression of *ybeX* in the absence of an inducer abolished the antibiotics sensitivity (**Fig. 3B**).

### The antibiotic sensitivity of ΔybeX depends on the growth history of cells

Our finding that while the *ΔybeX* cells have a lengthened lag phase during outgrowth from stationary phase, they appear to retain similar levels of metabolic activity during this lag phase to the WT cells, as well as similar exponential growth rate (**Fig. 1D**), led us to hypothesize that any cellular defects conferred by the lack of YbeX may accumulate during the late growth, preceding entry into the stationary phase and/or in the stationary phase itself. Such a stochastic process could lead to the observed single-cell level growth heterogeneity (**Fig. 2**). Accordingly, we assayed whether the phenotypes of *ΔybeZ*, *ΔybeY,* and *ΔybeX* depend on the growth phase where the cells originate. We surmised that if the *ΔybeX* phenotype is caused by a gradual accumulation of harm, then cells that have been given ample time to accumulate such harm, should exhibit a stronger phenotype than cells with only a few divisions.

First, we tested the antibiotic sensitivity phenotype. In this experimental setup, we start by growing a single bacterial colony for 12 hours into the early stationary phase (**Fig. 4A**). Then the experiment is divided into two. In the first arm, to assay the antibiotic sensitivity during outgrowth, stationary liquid cultures are directly dot spotted into agar plates containing sub-inhibitory concentrations of antibiotics. In the second arm, to assay the antibiotic sensitivity of exponentially growing cells, the same stationary cultures are first diluted a hundred-fold into fresh liquid media and grown at 37°C for four to five cell divisions until OD_600_ reaches 0.2-0.4, after which they are dot spotted.

**Fig. 4.**
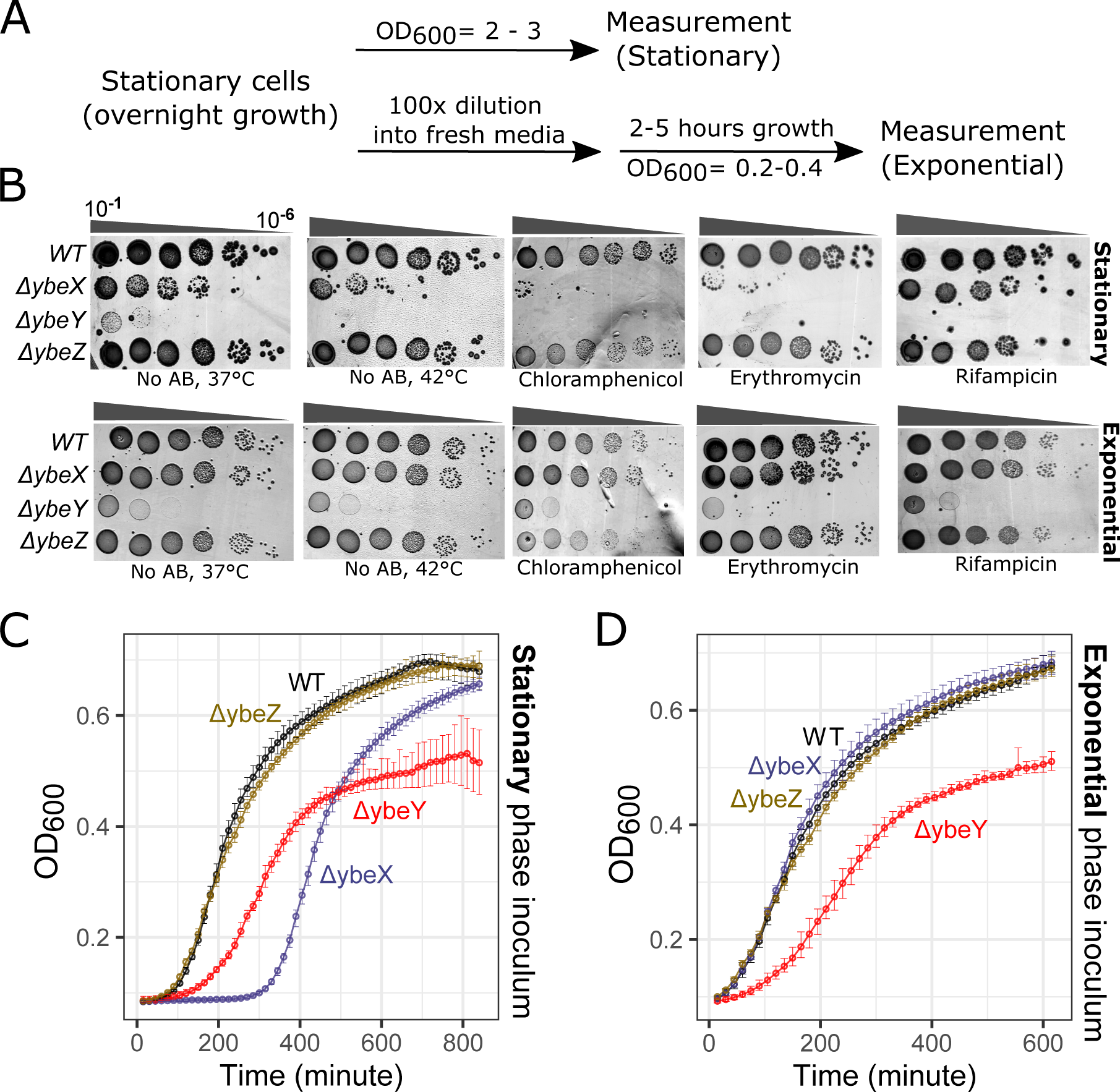
Growth phenotypes of *ΔybeX* are growth-phase dependent. (**A**) Experimental scheme for bacterial growth in LB medium. Overnight cultures were directly used for the Stationary phase experiments, while the cells were diluted into fresh LB and regrown for Exponential phase experiments. (**B**) Dot spot experiments of Keio wild-type (WT) and *ybeX*, *ybeZ*, and *ybeY* deletion strains. The plates were incubated at 37°C, except for the 42°C plate. (**C** and **D**) Growth curves of indicated *E. coli* strains grown on 96-well plates. The monitored growth of stationary (**C**) and exponential (**D**) phase cells of wild type and *ybeY*, *ybeZ*, and *ybeX* deletion strains in liquid LB medium at 37°C. The growth curves are presented as curves for four biological replicates from three independent experiments, where shaded areas represent the 95% CI-s.

The *ΔybeX* strain exhibited very strong chloramphenicol and erythromycin sensitivity in cells originating from early stationary phase, but no sensitivity to Rifampin (**Fig. 4B**). In contrast, the *ΔybeX* cells plated on the antibiotic after only a few rounds of the division had WT-like sensitivity to all tested antibiotics. In comparison, *ΔybeZ* cultures had similar intermediate levels of sensitivity to chloramphenicol, regardless of the growth history of cells, while they are not sensitive to erythromycin, rifampicin, and tetracycline (**Fig. 4B**). Exponentially growing *ΔybeZ* cells in MOPS minimal medium, supplemented with 0.3% glucose as the carbon source, also exhibited sensitivity to chloramphenicol (**Figure S4a**). *ΔybeY* cells had a very strong sensitivity to all tested antibiotics under both growth conditions. This is not surprising, considering its strong growth phenotype.

Testing the culture growth in liquid media, after diluting the culture directly from the early stationary phase, again showed a lengthened lag phase for *ΔybeX* but not for *ΔybeZ*, while the exponential growth rates of both *ΔybeX* and *ΔybeZ* were very similar to WT (**Fig. 4C**). The *ΔybeY* strain behaves similarly in both experiments, exhibiting a reduced exponential growth rate and reaching a lower maximal cell density. In contrast, when the cells are outgrown from exponential phase cultures, the WT, *ΔybeZ* and *ΔybeX* strains grow equally well, with no visible lag phase, while the *ΔybeY* strain has a reduced growth rate and a lower growth end-point, as expected (**Fig. 4D**).

### *ΔybeX* cells accumulate rRNA fragments

As *ybeX* is located in the same operon with *ybeY,* whose role is implied in ribosome assembly, we assessed the rRNA profiles of WT Keio and *ΔybeZ*, *ΔybeY*, and *ΔybeX* strains by formaldehyde denaturing agarose gel electrophoresis of total cellular RNA. In exponentially growing *ΔybeY* cells, we saw a substantial accumulation of immature 16S rRNA (17S rRNA), while *ΔybeX* and *ΔybeZ* cells had comparable levels of 17S rRNA to wild-type (**Fig. 5A**). *ΔybeY* cells also accumulate a faster-migrating 16S rRNA species, labelled as 16S* (**Figure 5A;** see also (Davies *et al*., 2010)). When we assessed the RNA extracted from stationary phase cultures in *ΔybeX* cells we observed a major RNA fragment of about a thousand nucleotides (**Fig 5A**). This fragment was not present in material obtained from exponentially grown *ΔybeX* cells. The wild type, the *ΔybeZ* and the *ΔybeY* cells exhibited no such fragments in either stationary or exponential cells.

**Fig. 5.**
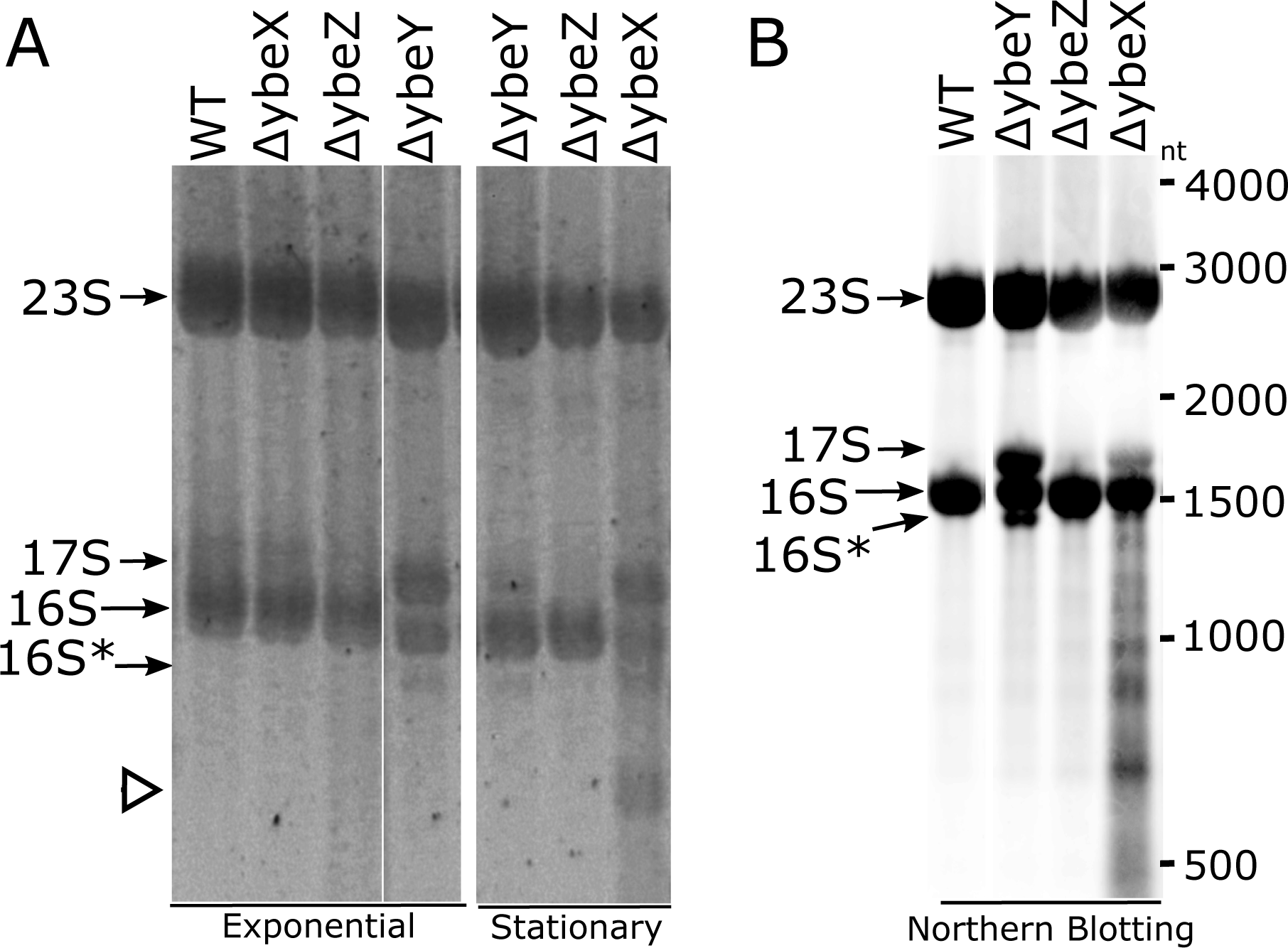
Accumulation of ribosomal RNA fragments in stationary phase *ΔybeX* cells. (**A**) Denaturing Agarose gel electrophoresis of hot phenol extracted total RNA samples from wild type and *ybeY*, *ybeZ*, and *ybeX* deletion strains. The empty triangle marks the accumulated shortened rRNA species. (**B**) Northern blot analysis of hot phenol extracted total RNA samples from wild type and *ybeY*, *ybeZ*, and *ybeX* deletion strains. The membrane was hybridized with 16S rRNA targeting oligonucleotide.

We used a more sensitive assay, the Northern blotting, on total RNA. **Fig. 5B** shows, for *ΔybeX* lysates, a wide spectrum of 16S rRNA intermediates ranging from 500 nt (our lower detection limit) to almost full length 16S rRNA. Note that due to apparent cross-binding of our 16S-targeting probe to the 23S rRNA, we also see the 23S rRNAs as distinct bands in the gel, but importantly there are no degradation fragments between the full length 23S rRNA and 17S rRNA in any of the strains. Also, the 17S pre-rRNA is present for *ΔybeX* and *ΔybeY*. Interestingly, *ΔybeX* does not contain the 16S* rRNA species, which is present in *ΔybeY* (but not *ΔybeX*). Except for the 16S* rRNA of *ΔybeY*, the WT, *ΔybeZ* and *ΔybeY* lanes lack degradation intermediates. Thus the ybeX cells contain a unique and disparate mixture of 16S rRNA degradation intermediates.

### The *ΔybeX* strain accumulates distinct rRNA species already during the late exponential growth

As there is neither assembly nor degradation of mature ribosomes in the early stationary phase (Piir *et al*., 2011), we conjectured that the 16S fragments observed in *ΔybeX* cells were likely accumulating by the late exponential phase. Accordingly, we purified, from late exponential cells, ribosomal subunits by sucrose gradient fractionation and analyzed the rRNA composition of the 70S ribosomes, as well as 50S and 30S subunits by Northern blotting. In these experiments we used probes specific for both ends of 17S precursor, for the 16S 3’ end, and for the mature 16S rRNA and 23S rRNA, allowing us to see degradation intermediates emanating from immature pre-16S rRNAs (**Fig. 6A**).

**Fig. 6.**
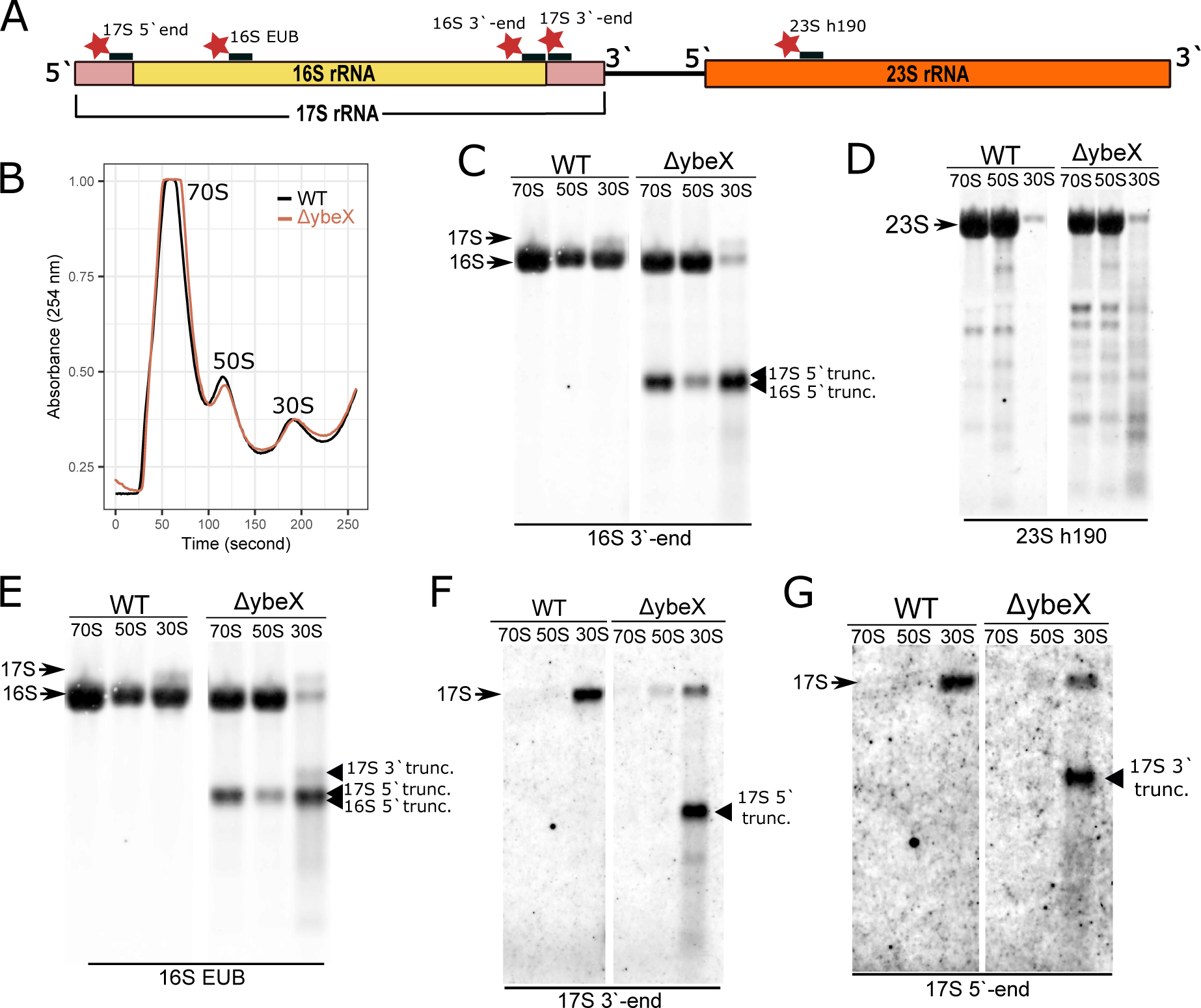
Deletion of *ybeX* leads to accumulation of distinct rRNA species. (**A**) rRNA operon illustration with locations of the Cyanine 5 (red star) labelled oligonucleotides. (**B**) Sucrose gradient profiles of WT and *ΔybeX* strains grown at 37°C for 2 hours after the OD600 reaches 0.3 (see Fig. 7A). 10-30% sucrose gradients were used for sedimentation. (**C**-**G**) Northern blot hybridization of the same membrane using different oligonucleotides labelled with Cy5. Truncated ribosomal RNA is annotated as “trunc.”.

The sucrose gradient profiles for WT and *ΔybeX* lysates are very similar, with the vast majority of ribosomal particles being in the presumably active 70S ribosome fraction and the small free subunit fractions exhibiting no obvious abnormalities (**Fig. 6B**). The Northern blots revealed 17S precursor rRNAs in the 30S fractions of both the WT and the *ΔybeX* strain, likely due to active ribosomal synthesis in both strains (**Fig. 6C, E**). In addition, in the ybeX strain the mature 16S rRNA species is substantially reduced in the 30S fraction, so that the 17S to 16S ratio is clearly shifted in relation to WT. Thus, in the *ΔybeX* strain the 30S fraction is unlikely to contain many functionally active ribosomal subunits.

In addition, there are two distinct 16S fragments, both around 1 kb long, in the ribosomal fractions originating from the *ΔybeX* cells. Firstly, there is a major 5’ end-truncated 16S rRNA fragment which is present in all ribosomal fractions, including the 70S ribosomes (**Fig. 6C,E**). In the 30S fraction this fragment is produced already from the 17S pre-rRNA (**Fig. 6F**), but the same fragment in the 70S ribosomes is not of this origin, presumably originating from full length mature 16S rRNA inside the 70S particles (**Fig. 6E**). Its presence in the 30S fraction is more pronounced than that of the 17S pre-rRNA, indicating that most of the pre-30S particles are inactive and degradation-bound in late exponential phase *ΔybeX* cultures. Secondly, there is a slightly larger 3’ end-truncated 16S RNA fragment (**Fig. 6E**), which is present in the 30S fraction only (**Fig. 6G**). This fragment also originates from the 17S precursor particles. In contrast, 23S rRNA specific probe reveals several relatively minor differences in degradation patterns between WT and *ΔybeX* strains (**Fig. 6D**).

Taken together, these results indicate that in the late exponential phase the majority of free 30S *ΔybeX* strain is in the process of being degraded. Moreover, the degradation fragments captured by the pre-16S rRNA specific probes strongly suggest that in the *ΔybeX* strain both pre-ribosomes (in the 30S fraction) and mature ribosomes (in the 70S fraction) are susceptible to degradation. While a majority of pre-ribosomes in the 30S fraction appear as the 1000-nt rRNA fragment (**Fig. 6E, F**), a minority of mature 70S is present as the 1000-nt fragment.

### *ΔybeX* perturbs ribosomal assembly through a separate mechanism from chloramphenicol

The strong sensitivity of *ΔybeX* cells to chloramphenicol (CAM) treatment prompted us to investigate the chloramphenicol phenotype further. CAM is a well-studied inhibitor of protein synthesis that binds to the large ribosomal subunit, inhibiting peptidyl transfer (Wilson, 2014). The effect of chloramphenicol on cell growth is at least partially mediated by the imbalanced synthesis of r-proteins, which results in the accumulation of partially assembled and misassembled ribosomal subunits (Siibak *et al*., 2009).

We tested the effect of sub-inhibitory concentrations of CAM on the ribosomes using sucrose gradient fractionation and northern blotting. Overnight-grown cells were diluted in liquid LB medium and grown until cells reached mid-exponential growth (OD_600_= 0.3). The CAM treatment took place for 2 hours. The cells were also grown without CAM for 2 hours as a control (**Fig. 7A**).

**Fig. 7.**
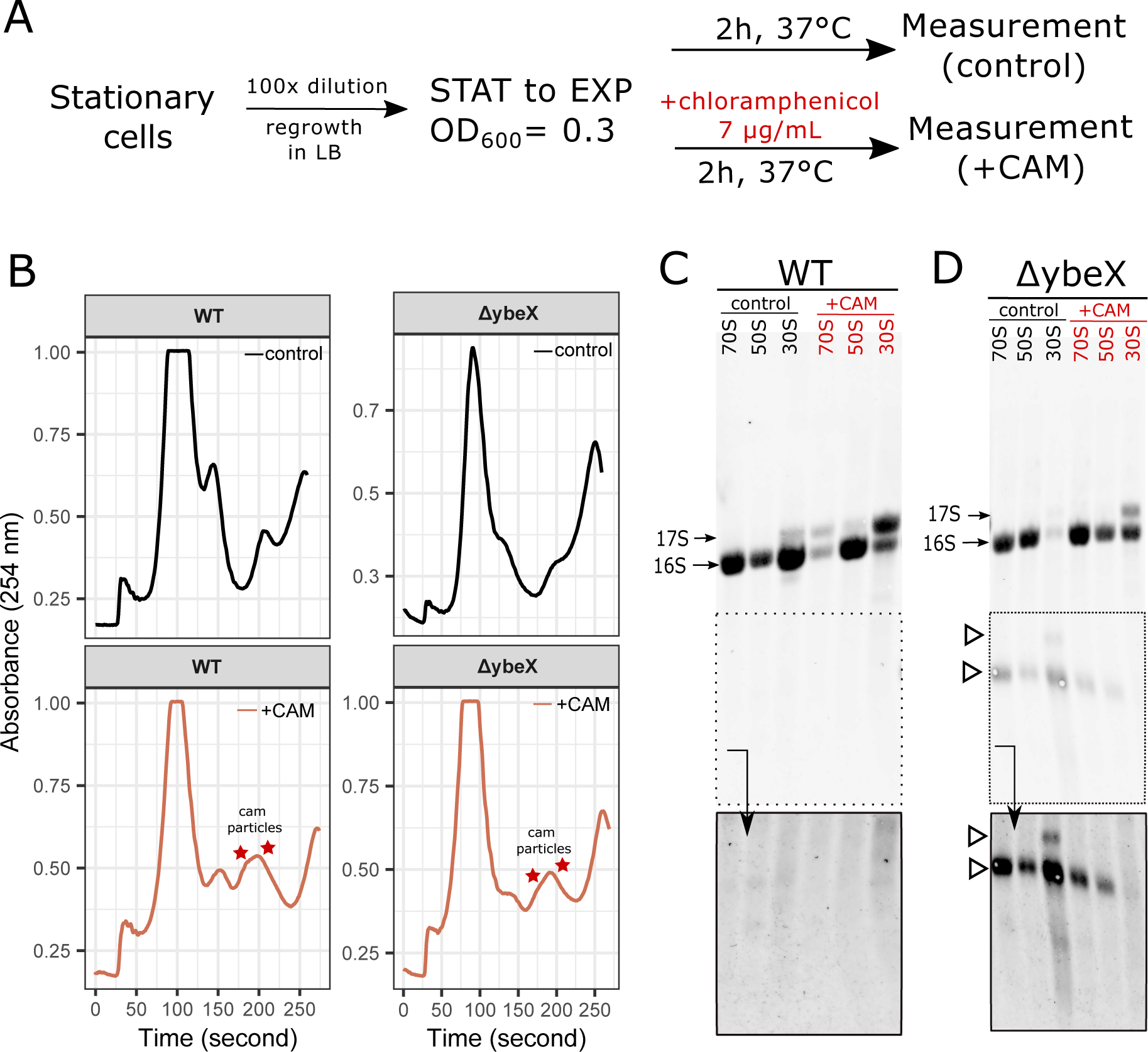
Accumulated distinct ribosomal RNA species are formed in vivo. (**A**) An experimental scheme where stationary phase cells (denoted as STAT) were grown to exponential phase (marked as EXP) followed by chloramphenicol (CAM) treatment (7µg/mL) for 2 hours. (**B**) 10-30% sucrose gradient fractionation of clarified WT and *ΔybeX* strains lysates. (**C**-**D**) Northern blot analysis of purified rRNA of sucrose gradient fractions separated on denaturing 1.5% agarose gel. The Northern blot was performed using 16S rRNA-specific oligo. The lower panels present the more prolonged exposure of the distinct accumulated rRNA species for more precise visualization.

While the CAM particles were formed in both wild type and *ΔybeX* strains (**Fig. 7B**), we failed to observe any aberrant rRNA species for the WT strain (**Fig. 7C**), while the accumulation of the distinct rRNA species appeared in *ΔybeX* cells repeatedly (**Fig. 7D**). Thus, the mechanism that leads to the degradation of pre-rRNA in 30S particles in *ΔybeX* cells seems to be different from that of the imbalanced protein synthesis caused by CAM. The CAM action mechanism also appears to stabilize *ΔybeX* 30S particles, while the pre-16S rRNA degradation intermediate is present in both 70S and 50S fractions. We believe its presence in the 50S to be due to cross-contamination from the 70S fraction. Interestingly, while WT CAM 70S particles contain a good measure of 17S pre-rRNA (which is absent in WT non-treated cultures), the *ΔybeX* CAM 70S particles, although containing the degradation intermediate, do not have this pre-16S rRNA species. These results suggest that the perturbation of assembly by CAM and by *ΔybeX* go by separate and at least partially independent mechanisms.

### The *ΔybeX* phenotype can be suppressed by MgCl_2_

As YbeX has been implicated in Mg^+2^ efflux (Gibson *et al*., 1991), we tested whether the supplementation of growth media with magnesium chloride affects the *ΔybeX* phenotype. First, we compared the growth of WT and *ΔybeX* in LB medium with and without magnesium supplementation (**Fig. 8A, Fig. S5a**). When the LB medium was supplemented with 10 mM MgCl_2_, the antibiotic sensitivity and heat shock phenotypes of *ΔybeX* disappeared (**Fig. 8A**). To test whether the effect is media-dependent, we used the SOB medium, which contains a high 10 mM concentration of MgCl_2_. Again, the phenotypes of *ΔybeX* disappeared (**Fig. S5a,b**). Thus, excess magnesium in the growth media, either LB or SOB, fully rescues the outgrowth growth phenotypes of the *ΔybeX* cells.

**Fig. 8.**
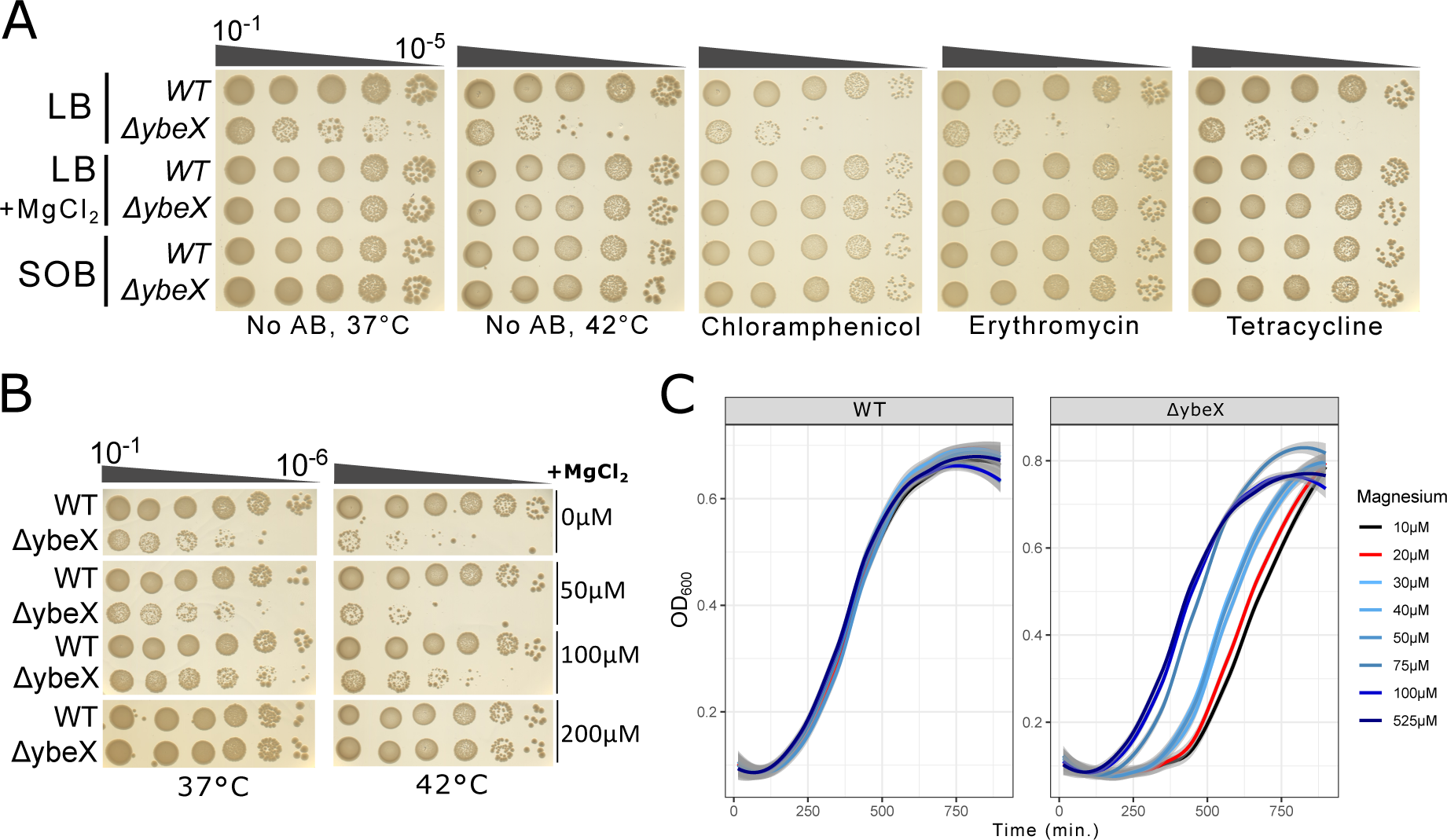
Magnesium supplementation rescues the *ΔybeX* phenotypes in various growth media. (**A**) The WT and *ΔybeX* cells were grown overnight in LB, SOB growth media, or LB supplemented with 10 mM MgCl_2_. The cells were serially diluted and spotted on LB Agar plates. The plates with antibiotics were incubated at 37°C. There are also controls without antibiotics, denoted “no AB”, at 37°C and at 42°C, as indicated in the first two sub-panels. (**B**) A single colony of WT or *ΔybeX* was grown overnight in the magnesium-limited peptide-based medium. PBM, 0 μM denotes no MgCl_2_ supplementation; otherwise, PBM is supplemented with 50, 100 and 200 μM MgCl_2_. (**C**) WT or *ΔybeX* cells were grown overnight in defined MOPS minimal media supplemented with indicated concentrations of MgCl_2_ and 0.3% glucose as carbon source. The outgrowth from these stationary phase cultures was done in MOPS, supplemented with 525 μM of MgCl_2_ (this is the prescribed optimal magnesium concentration of the 1x MOPS minimal media). The regrowth of the cells was monitored using a 96-well plate reader. The growth curves are presented as LOESS curves for six biological replicates from three independent experiments, where shaded areas represent the 95% CI-s for the fitted LOESS curve.

To test whether magnesium-deficient-rich media could increase the severity of the growth phenotype, we used the peptide-based medium (PBM), a rich, magnesium-limited, buffered, complex growth medium (Christensen *et al*., 2017). We used the PBM because it is free of any cell extract, which is the primary source of magnesium in almost all complex media (Li *et al*., 2020). To avoid diauxic inhibition, we modified it to contain casamino acids instead of glucose as the carbon source ((Blaiseau and Holmes, 2021); see Materials and Methods). When PBM is not supplemented with any MgCl_2_, the heat shock phenotype at 42°C is somewhat more substantial than in LB medium(**Fig. 8B**, compare **Fig. S5c with Fig. S5d**). Supplementing PBM with 50 μM and 100 μM MgCl_2_ partially suppresses the phenotype, first in 37°C and then in 42°C, and supplementation with 200 μM MgCl_2_ completely suppresses it under both temperature conditions (**Fig. 8B**). Therefore, we have a potentially sensitive regulatable system for driving the growth phenotype of the *ΔybeX* strain in a quantitative manner.

To test the sensitivity and robustness of such an experimental system, we did a liquid medium growth experiment in defined MOPS minimal medium, with glucose as the carbon source. Unlike with the PBM, in the MOPS medium, we can precisely control the magnesium levels by adding MgCl_2_ from 10 μM to 525 μM (the “normal” optimal level for this medium; (Christensen *et al*., 2017)). When the WT cells grew into an overnight stationary phase in different Mg^2+^-depleted MOPS media, there was no Mg^2+^ supplementation effect for the outgrowth lag phase duration (**Fig. 8C**, the left sub-panel). As expected, there was no effect on the actual growth rate after the lag phase. Under the same conditions, the Mg-supplementation effect on *ΔybeX* lag phase was very different (**Fig. 8C**, the right sub-panel). There appears a threshold effect, whereby Mg^2+^-supplementations by 50 μM and less produce a small gradual shortening of the lag phase from around 400 minutes to 350 minutes, while supplementation with 75 μM MgCl_2_ suddenly shifts the lag time to about 200 minutes, after which additional magnesium has little effect on the duration of the lag phase.

## DISCUSSION

In this work, we show that the putative Co^2+^/Mg^2+^ efflux protein YbeX is functionally involved in ribosome metabolism in *Escherichia coli*. We propose that during growth in the absence of *ybeX* there is a gradual accumulation of harm, possibly consisting of pre-17S rRNA and 16S rRNA partial degradation products (**Fig. 6C-G**), which will necessitate a longer lag phase upon outgrowth in a fresh medium. During this prolonged lag phase, the *ΔybeX* cells are metabolically active (**Fig. 1D**) and would be busy cleaning up the partial rRNAs, before new ribosome synthesis and subsequent cell division can commence. Interestingly, when we prolong the incubation of stationary phase cultures from overnight to 48 hours using MOPS minimal medium agar plates, the *ΔybeX* outgrowth phenotype and AB sensitivity disappear (data not shown). These findings suggest that the harm, which has been gradually accumulated, can also be gradually removed, in stationary cells with lower metabolic activity (**Fig. S1f, Fig. S2**). Intriguingly, although the mid-to late-exponential phase *ΔybeX* cells accumulate rRNA degradation products, to some extent even in the 70S fraction containing mature translation-ready ribosomes (**Fig. 6C,E**), they have very much WT-like sucrose gradient profiles (**Fig. 6B**), indicating no accumulation of major ribosome-like particles. In addition, although the *ΔybeX* cells have a clear growth phenotype, manifesting itself in a lengthened outgrowth lag phase and sensitivity to antibiotics, the exponential growth rate of the *ΔybeX* cells is indistinguishable from WT, as are the growth end-points (**Fig. 1C, Fig. 8C**).

The growth phenotype of *ΔybeX* is Mg^2+^ dependent, being present only in Mg^2+^-limiting growth conditions (**Fig. 8**). Intriguingly, the rescue of *ΔybeX* growth by MgCl_2_ occurs through a threshold effect, whereby something happens between 50 µM and 75 µM MgCl_2_ that abolishes the phenotype essentially in one fell swoop. This suggests that Mg^2+^-deprivation does not lead to the growth defect through a thousand small pricks, i.e. through many individual Mg^2+^-binding sites of different strength and metabolic functions, but more likely through a mechanism that quickly becomes saturated between 50 µM and 75 µM of added MgCl_2_. As most Mg^2+^ ions in the cell bind to the ribosome (>170 Mg^2+^ per ribosome, up to 70 000 ribosomes per cell) (Akanuma, 2021) and there are about a hundred Mg^2+^ions that specifically bind to one or two non-bridging phosphate oxygen’s of the large subunit rRNA (Klein *et al*., 2004), it is tempting to speculate that the observed magnesium threshold corresponds to activation of the ribosome by structural stabilization. Intriguingly, and in accordance with the role of YbeX in maintaining strict Mg^2+^-homeostasis, we found that overexpression of the *ybeX* from a high-copy plasmid, is toxic to both WT and *ΔybeX* cells, even in the absence of inducer.

What could be the mechanism of action of the YbeX protein on the ribosome? Unlike its neighboring gene products, the YbeY and the YbeZ, there is no evidence that YbeX binds to the ribosome or any ribosome-associated protein. Nonetheless, at this stage, we cannot exclude the possibility of a direct action of the YbeX on the ribosome. The *ybeX*/*corC* gene was initially recovered in *Salmonella typhimurium* in a screen for resistance to cobalt and proposed to contribute, possibly as a co-effector of CorA, to the efflux of divalent cations (Gibson *et al*., 1991). As yet, there is no mechanistic function ascribed to this gene, and while Mg^2+^ influx is generally well-studied, its efflux is poorly understood in bacteria (Armitano *et al*., 2016). Essentially, YbeX is a cytoplasmic protein (Sueki *et al*., 2020), for which we have indirect evidence that it might be somehow involved in Mg^2+^-efflux. Our finding that the growth phenotype of the *ΔybeX* strain requires low extracellular Mg^2+^ is consistent with the role of YbeX in Mg-efflux, as Mg^2+^ efflux is known to be inhibited at low extracellular magnesium (Nelson and Kennedy, 1971) and can be activated by adding 1mM MgCl_2_ to the growth medium for *S. typhimurium* (Gibson *et al*., 1991).

YbeX has been genetically connected to translation, as *E. coli* cells, relying for growth on an artificial ribosome variant, where the subunits are covalently tethered by fused rRNAs, need a nonsense mutation in the *ybeX* gene, together with a missense mutation in the *rpsA*, for faster growth (Orelle *et al*., 2015). In this case, a likely mechanism of action is that increased Mg^2+^ concentration in these cells stabilizes the artificial ribosomal construct and thus activates it for protein synthesis.

Thus, we currently favour the model whereby the effect of *ybeX* deletion on ribosomal metabolism is indirect, happening through an increased concentration of intracellular Mg^2+^. As both very low and very high Mg^2+^ concentrations are detrimental to cells, mainly through translation, the intracellular free Mg^2+^ is tightly controlled between 1 mM and 5 mM (Akanuma, 2021). *In vitro* translation is very sensitive to increased Mg^2+^ concentration, being >95% inhibited already at 6 mM MgCl_2_ (Borg and Ehrenberg, 2015), while 10 mM MgCl_2_ is enough to reduce translational accuracy by order of magnitude (Rodnina *et al*., 1999).

We also need to consider the effect of low extracellular Mg^2+^ on cell physiology. Here the Mg^2+^ acts as a counterion to neutralize the phosphate groups of outer-membrane lipopolysaccharides (Groisman and Chan, 2021). In addition, extracellular Mg^2+^ binds to many membrane proteins, stabilizing their structures. Accordingly, a lack of extracellular Mg^2+^ leads to permealization of the outer membrane, including to hydrophobic antibiotics like Erythromycin and Rifampin (Vaara, 1992).

Our current provisional model, whereby elevated intracellular free Mg^2+^ leads to growth defect through promoting ribosomal degradation, differs from the one worked out in Eduardo Groisman’s lab (Pontes *et al*., 2016). The Groisman model proposes an active role for the ribosome in magnesium homeostasis, mainly under low concentrations of intracellular free Mg^2+^. Specifically, low cytosolic Mg^2+^ causes, in a genetically controlled manner, a shut-down in ribosome production and a reduction in the cellular ATP levels, which in turn leads to reduced proteolysis by ATP-dependent proteases and to reduced protein synthesis (Groisman and Chan, 2021). This Mg^2+^-dependent control supersedes rRNA transcriptional control by amino acid availability and ATP abundance. Unifying these models into a more general view of the roles of Mg^2+^ in ribosomal metabolism requires more research. Nevertheless, we can safely say that the role of magnesium homeostasis in ribosomal metabolism is becoming an increasingly fertile field of study.

## MATERIALS AND METHODS

### Bacterial Strains, Plasmids and Growth Media

Genotypes of bacterial strains, plasmid descriptions, and sequences of primers used in this study are listed in Tables S2-S3. Bacteria were grown in Difco^TM^ LB Broth (BD brand #240230 consist of Tryptone 10 g/L, Yeast Extract 5 g/L, Sodium Chloride 5 g/L). LB agar plates were prepared from Difco^TM^ LB Agar (BD brand #240110). Antibiotics used were Ampicillin (100 μg/mL) or Carbenicillin (100 μg/mL), Chloramphenicol (25 μg/mL), Kanamycin (50 μg/mL), andTetracycline (12.5 μg/mL).

Keio collection deletion strains including *ΔybeX*, *ΔybeY*, and *ΔybeZ*, and *Escherichia coli* wild-type BW25113 strains were used in this study (Baba *et al*., 2006). We also reconstructed the *ybeX* single deletion strain using *E. coli* MG1655 and BW25113 via lambda red recombination (Datsenko and Wanner, 2000). The removal of kanamycin resistance gene (*kan*) from the bacterial chromosome was performed using the pCP20 plasmid.

*E. coli DH5α* strain was used for plasmid cloning and propagation. The TransBac library, a new

*E. coli* overexpression library based on a single-copy vector, was obtained from Dr. Hirotada Mori (Nara Institute of Science and Technology, Japan) as a stab stock (Otsuka *et al*., 2015).

### Conjugation of the TransBac library plasmids

*Hfr* strain is the donor strain that carry each TranBac library plasmid and capable of transfer of the target plasmid by conjugation (unpublished resource by Mori). The donor strain was grown on LB agar plates supplemented with Tetracycline (Tc) and 25 µg/mL diaminopimelic acid (DAP). *Hfr* strain requires DAP because of the deletion of *dapA* gene. Well grown donor and acceptor cell cultures were mixed 1:1 ratio in a 1.5 mL polypropylene tube, and incubated at 37°C for 1 hour without shaking. After appropriate time for conjugation, the cell mix was plated onto LB agar plates containing Tc (12.5 µg/ml) without DAP. The plates were incubated at 37°C overnight.

### Construction of the TransBac empty (pTB-empty) plasmid

The single-copy TransBac library plasmid coding *ybeX* was purified using in-house alkaline lysis method followed by purification via FavorPrep™ plasmid DNA extraction mini kit (Favorgen, Austria). The cloning site was sequenced by Sanger sequencing. The *ybeX* coding region was removed via restriction enzyme cleavage of *XmaJI* and *SfiI* (Thermo Scientific™). The sticky ends were filled using Klenow fragment (Thermo Scientific™) and the linear plasmid was ligated using T4 DNA ligase (Thermo Scientific™) following manufacturer protocols. The ligation reaction was transformed into Inoue *E. coli DH5α* chemical competent cells (Green and Sambrook, 2020), and the TransBac empty backbone plasmid was purified as mentioned above. The size of the plasmid DNA was determined via agarose gel electrophoresis, and the cloning site was sequenced. The plasmid was electroporated into Keio collection strains BW25113 and *ΔybeX*.

### Preparation of the Peptide Based Media (PBM)

Growth in peptide-based media (PBM) is magnesium-limited (Christensen *et al*., 2017). The previously described PBM recipe requires addition of 0.4% of glucose (4 g/L) as carbon source. We prepared the PBM via dissolving 10g/L Peptone, 1.5% casein hydrolysate (casamino acids) and 40 mM MOPS (3-(N-morpholino) propane sulfonic acid) buffer pH 7.4. The 50x MOPS buffer stock solution contained 2M MOPS and 0.2M Tricine pH 7.4 set with concentrated KOH.

We achieved extremely high cell densities, OD_600_ = 10-13, when 20 g/L peptone, 3% Casein hydrolysate were used in the presence of 1-10 mM MgCl_2_ or MgSO_4_.

### Bacterial Spot Assay

Bacterial cell cultures were diluted to final OD_600_ = 0.125 which was the first dilution (10^-1^), and then 10x serial dilutions were applied. 5µL of each dilution were spotted on LB Agar plates with or without antibiotics (No AB) supplementation. The plates were imaged using an Epson Expression 1680-pro scanner.

### Colony Size Characterization and Quantification

Keio wild-type and *ΔybeX* strains were grown overnight in LB or defined MOPS minimal media (Neidhardt *et al*., 1974). Well-grown bacterial cell cultures were serially diluted and plated on LB agar plates using glass beads (Hecht Assistent, #41401004). We aimed to have approximately 100 colonies per plate. The plates were incubated overnight at 37°C or 42°C and scanned using EPSON Expression 1680pro scanner. The images were subjected to AutoCellSeg software (Khan *et al*., 2018). The colonies were first picked automatically using program default settings and then, as a second step, manual picking was applied (picking small colonies, deselecting adherent colonies, etc.). The data was analyzed in R::tidyverse package (Wickham *et al*., 2019; R Core Team, 2022).

### Growth monitoring in the 96-well plate reader

*C*ells were diluted in the appropriate growth media to OD_600_ = 0.55, and 10 µL of the diluted cells were transferred into 100 µL of growth media in a 96-well plate. The 96-well plate edges were filled with distilled water. The remaining 60 wells were used to monitor the growth. At least one column was always set as negative control. Alamar Blue reagent (BioRad, #BUF012B) was used according to the manufacturer protocol.

### Sucrose gradient fractionation

*E. coli* strains from Keio collection were streaked onto LB agar plates and grown overnight at 37°C. A single colony of each strain was inoculated into LB and aerated at 37°C overnight. In the following morning, the culture densities were determined via spectrophotometer (Biochrom Ultrospec 7000), the cells were diluted to a final OD_600_ of 0.05-0.06 in LB medium (150-250mL) and grown until OD_600_ = 0.3-0.35. The cultures were then split into two flasks, in which the chloramphenicol treatment was carried out, while the other was grown as a control for 2 hours.

The cells were transferred into centrifugation bottles, cooled on ice and pelleted at 4000xg, at +4°C for 10 minutes. The supernatant was removed, and the cell pellet was snap-frozen in liquid nitrogen and stored at −80°C. The cells were dissolved in 1 mL of lysis buffer consist of 25 mM Tris-HCl pH 7.9, 60 mM KCl, 60 mM NH_4_Cl, 6 mM MgCl_2_, 5% glycerol supplemented with 1mM PMSF, protease inhibitor (Roche, #04693159001) and 5mM βME added freshly to the buffer before the lysis. The cells were lysed using FastPrep homogenizer (MP Biomedicals) by three 40-second pulses at 4.0 m/s with chilling on ice for 5 min between the cycles. The beads were purchased from BioSpec Products, and 0.4 gram of 0.5mm Zirconia/Silica beads (BioSpec, #11079105z) and 0.9 gram of 0.1mm Zirconia/Silica beads (BioSpec, #11079101z) was used.

The lysate was clarified by centrifugation 16,100xg for 40 minutes at 4°C. Clarified lysates were treated with 50 units/ml DNase I (MN, #740963). The lysates were loaded onto 10-30% sucrose gradients in a buffer containing 25mM Tris-HCl pH 7.9, 100mM KCl, 10mM MgCl_2_, supplemented with 5mM βME. The gradients were centrifugated at 20,400 rpm for 17 h at 4°C in a SW-28 Beckman Coulter rotor (ω^2^t=2.8e+11). The samples from the gradient were pumped starting from the bottom through a spectrophotometer (Econo UV Monitor, BIO-RAD), which can detect A254 as a readout. The data was recorded by Data Acquisition software (DataQ Instruments) and imported into R for plotting (R Core Team, 2022).

### Purification of rRNA from Ribonucleoprotein (RNP) Complexes

Ribosomes and ribosomal subunits were collected from sucrose gradients as peak fractions. The sucrose fractions were collected into 15 mL falcon tubes and diluted by at least two-fold with the gradient buffer (25mM Tris-HCl pH 7.9, 100mM KCl, 10mM MgCl_2_). Next, 2.5 vol. of 96% ethanol was added to the samples and incubated at −20°C overnight. The fractions were pelleted via centrifugation for 45 minutes at 4000 rpm +4°C. The pellet was washed with 70% EtOH and centrifugation was re-applied for 10 minutes. The ribonucleoprotein complexes were suspended in 0.1 mL of MilliQ water, and samples were stored at −20°C.

The rRNA was purified with phenol-chloroform extraction. The samples were kept on ice and 1% SDS-containing phenol was added to the samples. Samples were vortexed vigorously for 10 s, kept on ice for 5 min, and centrifuged at 16,200xg at +4°C. The water phase was transferred to a new microfuge tube into which chloroform:phenol mixture (1:1) was added and vortexed for 10 seconds. This step was repeated, using only chloroform to avoid phenol carryover. The water phase was transferred to a new microfuge tube, and the RNA was precipitated with 2.5 vol. ethanol at −20°C for 1 hour. The pellet was washed with 70% EtOH and dried at room temperature for 5 minutes. The purified RNA was dissolved in ultra-pure distilled water.

### Total RNA Purification using hot phenol extraction

The strains were grown in LB at 37°C. 10-12 mL of cell culture were transferred to a 15 mL Falcon tube, pelleted via centrifugation at 8000xg for 3-4 minutes, snap-frozen in liquid nitrogen, and stored at −80°C until RNA purification. Total RNA was purified with hot phenol-chloroform extraction, as described previously (Kasari *et al*., 2013).

### Denaturing Agarose Gel Electrophoresis

The isolated RNA samples were separated on denaturing 1.5% agarose gel containing 1xMOPS buffer and 2% formaldehyde. 5 μg of RNA (no more than 6.6 μL in final volume) was mixed with 5.4 μL of formaldehyde, 3 μL of 10x MOPS buffer and 15 μL of formamide. The samples and RNA markers from Thermo Scientific (RiboRuler High Range, #SM1821 and Low Range RNA ladder, # SM1831) were denatured at 55°C for 15 minutes. The RNA mixes were then cooled on ice. Sample loading dye (5μL, 1:6) (0.25% bromophenol blue, 40% sucrose) was added to the samples, and the samples were loaded onto the gel. The electrophoresis buffer was the same as the buffer used to prepare the gel, 1 x MOPS. During the first hour, 60V was applied, and then the voltage was increased to 85V.

After 5 hours, when the run ended, the ladder region was cut off and stained for 30 min in the running buffer containing 10000x diluted Diamond™ Nucleic acid dye (Promega). The transfer of the RNA from the agarose gel to the nylon membrane (Amersham Hyband™-N+, GE Healthcare, #RPN303B) was done as via capillary transfer of RNA from the denaturing agarose gel to the nylon membrane (Sambrook, 2001). UV crosslinking was applied to achieve RNA crosslinking to the nylon membrane.

### Hybridization of the Northern Blot Membrane

20-25 mL hybridization buffer (0.5 M Sodium phosphate buffer pH 7.2 containing 7% SDS) and the rotating bottle were heated in a hybridization oven (Hybrigene, #Z649570) at 62°C in darkness. The membrane was placed in the bottle and rotated for two hours. The fluorescent-labelled DNA oligonucleotide was added to 10 μM final concentration, and hybridization occurred overnight. Next day, the wash buffer (20 mM sodium phosphate buffer pH 7.2 containing 1% SDS) was warmed in a water bath to 43°C. The membrane was washed with this pre-warmed buffer in a temperature-controlled orbital shaker in a metal box, preventing light exposure. The membrane was washed thrice for 5 minutes with approximately 250 mL of the wash buffer at 43°C. Finally, the membrane was placed into a plastic envelope. The scanning of the membrane was done in the Amersham Typhoon™ laser scanner.

### Statistical analysis

Two-sided Student’s t test with unequal variances was done in GraphPad Prism version 5.01 (Graphpad Software, San Diego, CA, USA). Other statistical analyses were done in R vers. 4.2.1, using the Brms package v. 2-18-0 for Bayesian modelling (Bürkner, 2018). For joint multilevel modelling of mean colony radiuses and their standard deviation, the model employing Student’s t likelihood was, in brms model language, brm(bf(Radius∼Strain*temp + (1|day) + (1|plate), sigma∼Strain*temp + (1|day)+ (1|plate)), data=full, family = student(), prior= c(prior(normal(0, 5), class=b), prior(normal(0,2), class=“sd”), prior(normal(0,2), class=“b”, dpar=“sigma”))). For mixture modelling of mean colony radiuses, the model description is brm(Radius∼Strain*temp + (1|day) + (1|plate), data=full, family = mixture(gaussian, gaussian)). The alamarBlue and corresponding OD_600_ measurements shown in Fig. 1 were modelled with splines using the blmss package version 1.1-8 (Umlauf *et al*., 2021). The modelling was done separately for each strain and for each condition (Alamar signal and OD_600_ signal). The model description is bamlss(value ∼ s(Time_min), family=“gaussian”). The growth curves in Fig. 8C were done using the LOESS smoother in the ggplot2 package v. 3.4.0 geom_smooth function (Wickham, 2016; Wickham *et al*., 2019).

## ACKNOWLEDGEMENTS

We are grateful to Niilo Kaldalu (University of Tartu) for his helpful discussions and help in writing the manuscript. Furthermore, we thank Dr Hirotada Mori (Nara Institute of Science and Technology, Japan) for sending us the TransBac plasmid library. Finally, we thank Aksel Soosaar and Viia Kõiv (University of Tartu) for their technical advice and support. This work was supported by funds from the European Regional Development Fund through the Centre of Excellence for Molecular Cell Technology and the Estonian Research Council (grant PRG335).

## AUTHOR CONTRIBUTIONS

ÜM, TT conceived the study. İS, ÜM, and TT designed the research. İS, SR, EA and AŽ conducted the experiments. İS, ÜM, and MP analyzed the data. İS prepared the figures and tables. İS and ÜM wrote the manuscript. All authors read and approved the manuscript.

---

**Table S1.** Minimal inhibitory concentrations (MICs, µg/ml) of various antibiotics for Keio wild-type BW25113 and *ΔybeX* strains. Mentioned *E. coli* cells were grown in LB liquid medium and the MICs were determined by the dilution method according to EUCAST. (The European Committee on Antimicrobial Susceptibility Testing, Clinical Breakpoints—Bacteria (version 13.0). *ΔybeX* cells showed two times lower MIC values to the ribosome-binding antibiotics: tetracycline, chloramphenicol, erythromycin, clindamycin and fusidic acid.

| Antibiotics | Mechanism | BW25113 | $\Delta ybeX$ |
| --- | --- | --- | --- |
| Amikacin | Ribosome binding | 4 to 8 | 4 to 8 |
| Ampicillin | β-lactam/Cell wall synthesis | 5 | 5 |
| Chloramphenicol | Macrolide/Ribosome binding | 8 | 4 |
| Ciprofloxacin | Fluoroquinolone, DNA replication | 0.015 | 0.015 |
| Clindamycin | Ribosome binding | 160 | 80 |
| Erythromycin | Ribosome binding | 100 | 50 |
| Fusidic Acid | Steroid/Ribosome binding | 512 | 256 |
| Kanamycin | Aminoglycoside/Ribosome binding | 10 | 10 |
| Mupirocin | Polyketide/ RNA, protein synthesis | 80 | 80 |
| Norfloxacin | Fluoroquinolone, DNA replication | 0.125 | 0.125 |
| Ofloxacin | Quinolone, DNA replication | 0.08 | 0.08 |
| Polymyxin B | Antimicrobial peptide, Cell wall binding | 0.313 | 0.313 |
| Rifampicin | inhibiting RNA polymerase | 10 | 10 |
| Streptomycin | Aminoglycoside/Ribosome binding | 8 | 8 |
| Tetracycline | Ribosome binding | 4 | 2 |
| Tobramycin | Aminoglycoside/Ribosome binding | 5 | 5 |

**Table S2.**
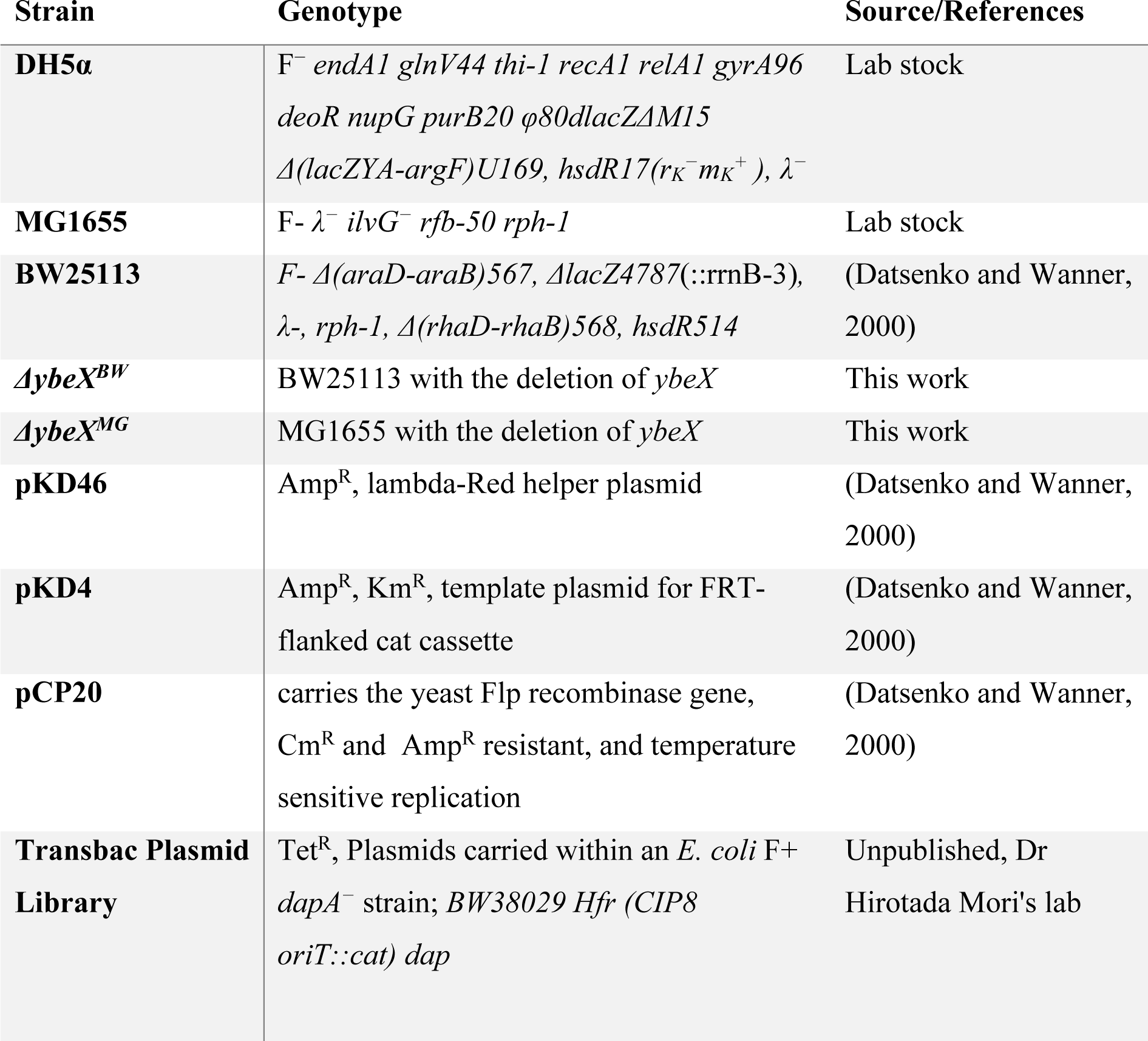
List of *E. coli* bacterial strains and plasmids used in this study. The abbreviations are - Cm^R^, chloramphenicol resistance; Amp^R^, ampicillin resistance; Km^R^, kanamycin resistance; Tet^R^, tetracycline resistance.

| Strain | Genotype | Source/References |
| --- | --- | --- |
| **DH5α** | F<sup>-</sup> *endA1 glnV44 thi-1 recA1 relA1 gyrA96 deoR nupG purB20 φ80dlacZΔM15 Δ(lacZYA-argF)U169, hsdR17(r<sub>K</sub><sup>-</sup>m<sub>K</sub><sup>+</sup>), λ<sup>-</sup>* | Lab stock |
| **MG1655** | F- λ<sup>-</sup> *ilvG<sup>-</sup> rfb-50 rph-1* | Lab stock |
| **BW25113** | F- Δ(*araD-araB*)567, Δ*lacZ*4787(::rrnB-3), λ<sup>-</sup>, *rph-1*, Δ(*rhaD-rhaB*)568, *hsdR514* | (Datsenko and Wanner, 2000) |
| **Δ*ybeX*<sup>BW</sup>** | BW25113 with the deletion of *ybeX* | This work |
| **Δ*ybeX*<sup>MG</sup>** | MG1655 with the deletion of *ybeX* | This work |
| **pKD46** | Amp<sup>R</sup>, lambda-Red helper plasmid | (Datsenko and Wanner, 2000) |
| **pKD4** | Amp<sup>R</sup>, Km<sup>R</sup>, template plasmid for FRT-flanked cat cassette | (Datsenko and Wanner, 2000) |
| **pCP20** | carries the yeast Flp recombinase gene, Cm<sup>R</sup> and Amp<sup>R</sup> resistant, and temperature sensitive replication | (Datsenko and Wanner, 2000) |
| **Transbac Plasmid Library** | Tet<sup>R</sup>, Plasmids carried within an *E. coli* F+ *dapA<sup>-</sup>* strain; BW38029 *Hfr (CIP8 oriT::cat) dap* | Unpublished, Dr Hirotada Mori's lab |

**Table S3.**
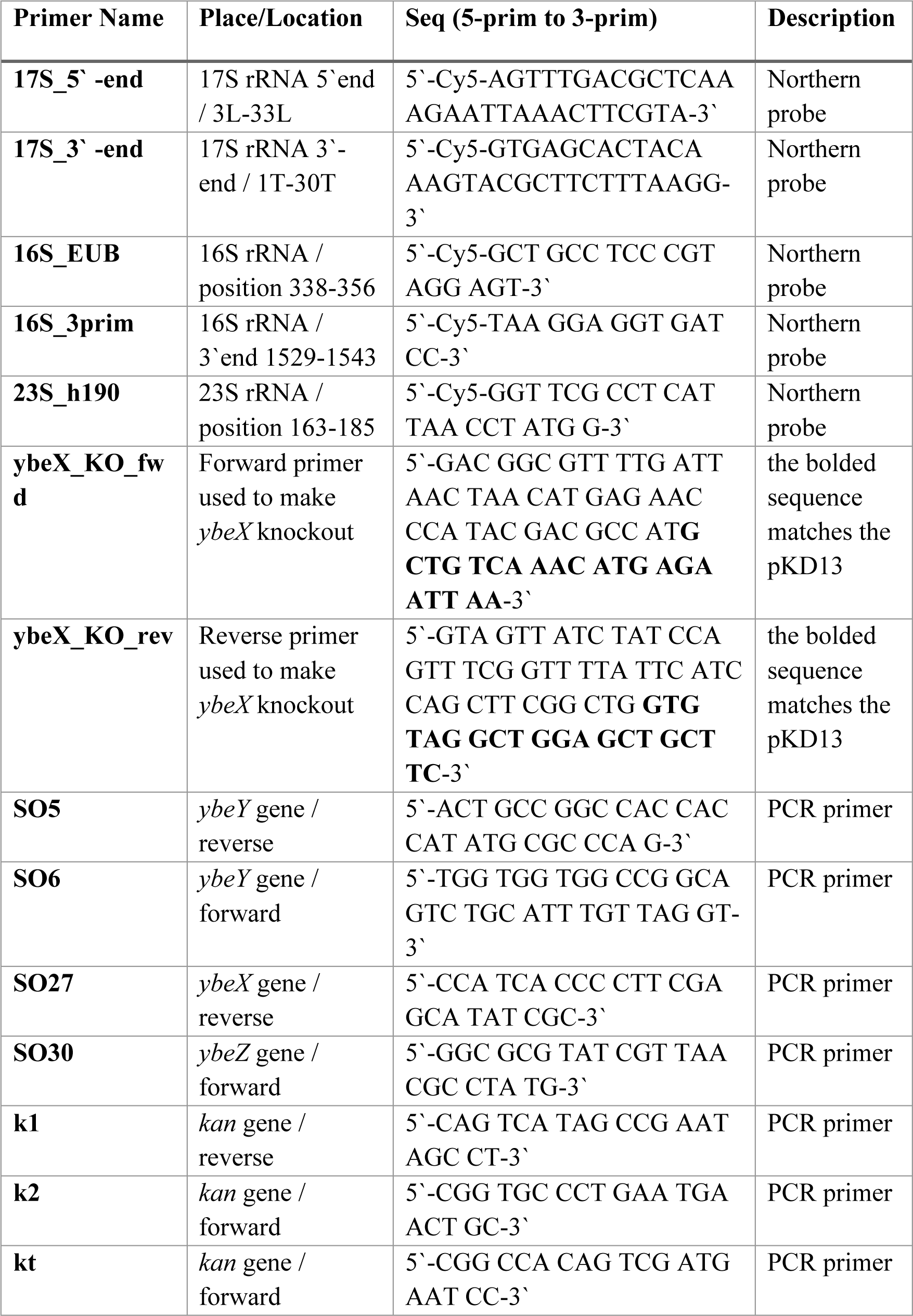
Oligonucleotides-Primers used in this study, including fluorescently (Cy5) labelled oligonucleotides. T-and L-stand for the trailer and leader tails of the 16S ribosomal RNA, respectively.

**Fig. S1.**
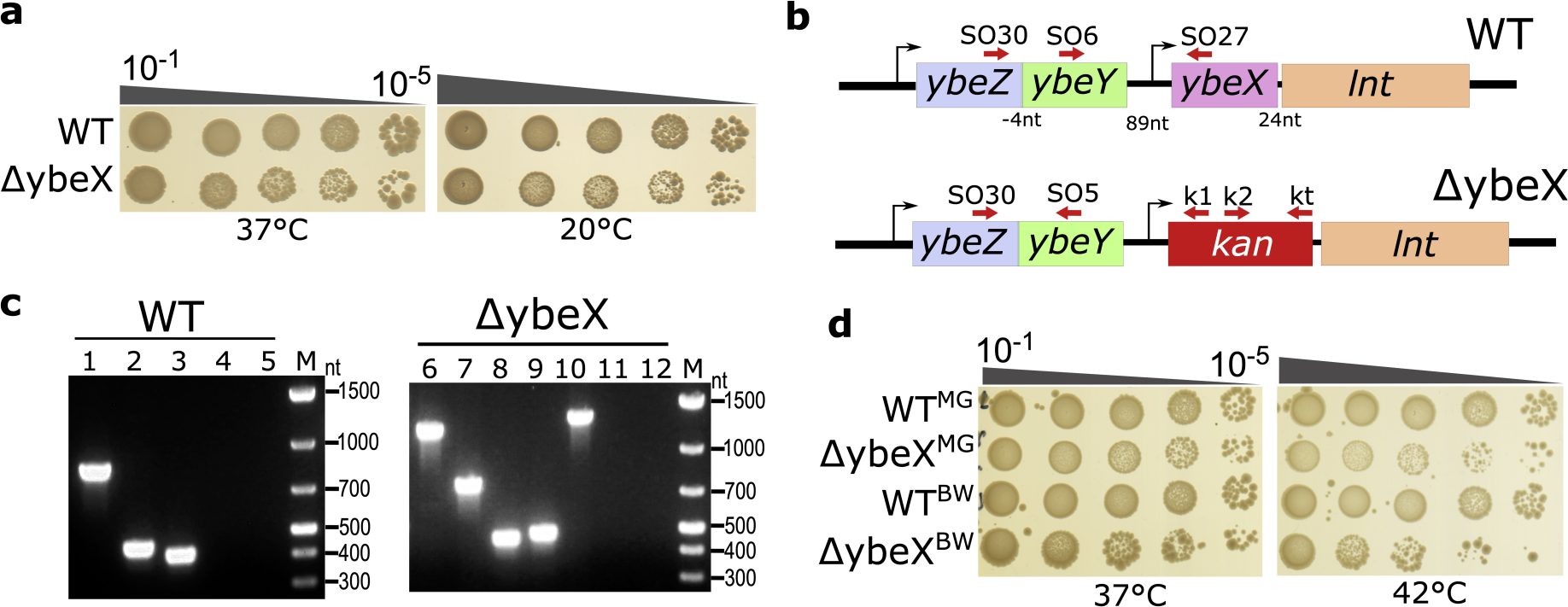
Deletion of *E. coli ybeX* leads to growth phenotypes. (**a**) Dot spot assay with wild-type and *ΔybeX* strain. The strains were grown in LB liquid medium overnight, diluted and spotted on LB agar plates. The agar plates were incubated at 37°C overnight or 20°C for 32 hours. (**b**) Chromosomal organization of the *ybeZYX-lnt* operon genes of wild type and *ΔybeX* in *E. coli*. The *ybeX* gene has been replaced with a kanamycin-resistant cassette (*kan*) in *ΔybeX*. Primers for colony PCR are indicated as red arrows. (**c**) Each band on the 1.2% agarose gel corresponds to a colony PCR reaction with different sets of forward and reverse primers 1-(SO30-SO27), 2-(SO30-SO5), 3-(SO6-SO27), 4-(SO6-k1), 5-(SO30-k1), 6-(SO30-k1), 7-(SO6-k1), 8-(SO30-SO5), 9-(k2-kt), 10-(SO6-kt), 11-(SO30-SO27), 12-(SO6-SO27), M-Thermo Scientific GeneRuler 1kb plus DNA ladder. The agarose gel contains 0.6 µg/mL ethidium bromide. Ethidium bromide wasn’t added to the running buffer. Negative controls were set for WT (empty lines 4 and 5) via primers aligned with the *kan* resistance gene. The primer SO27 don’t have the specific sequence to bind in *ΔybeX* (empty lines 11 and 12). (**d**) Dot spot assay for *E. coli* wild type MG1655 (WT^MG^) and Keio wild type BW25113 (WT^BW^) strains and respective chromosomal deletion of *ybeX* gene using lambda red homologous recombination (see Materials and Methods).

**Fig. S2.**
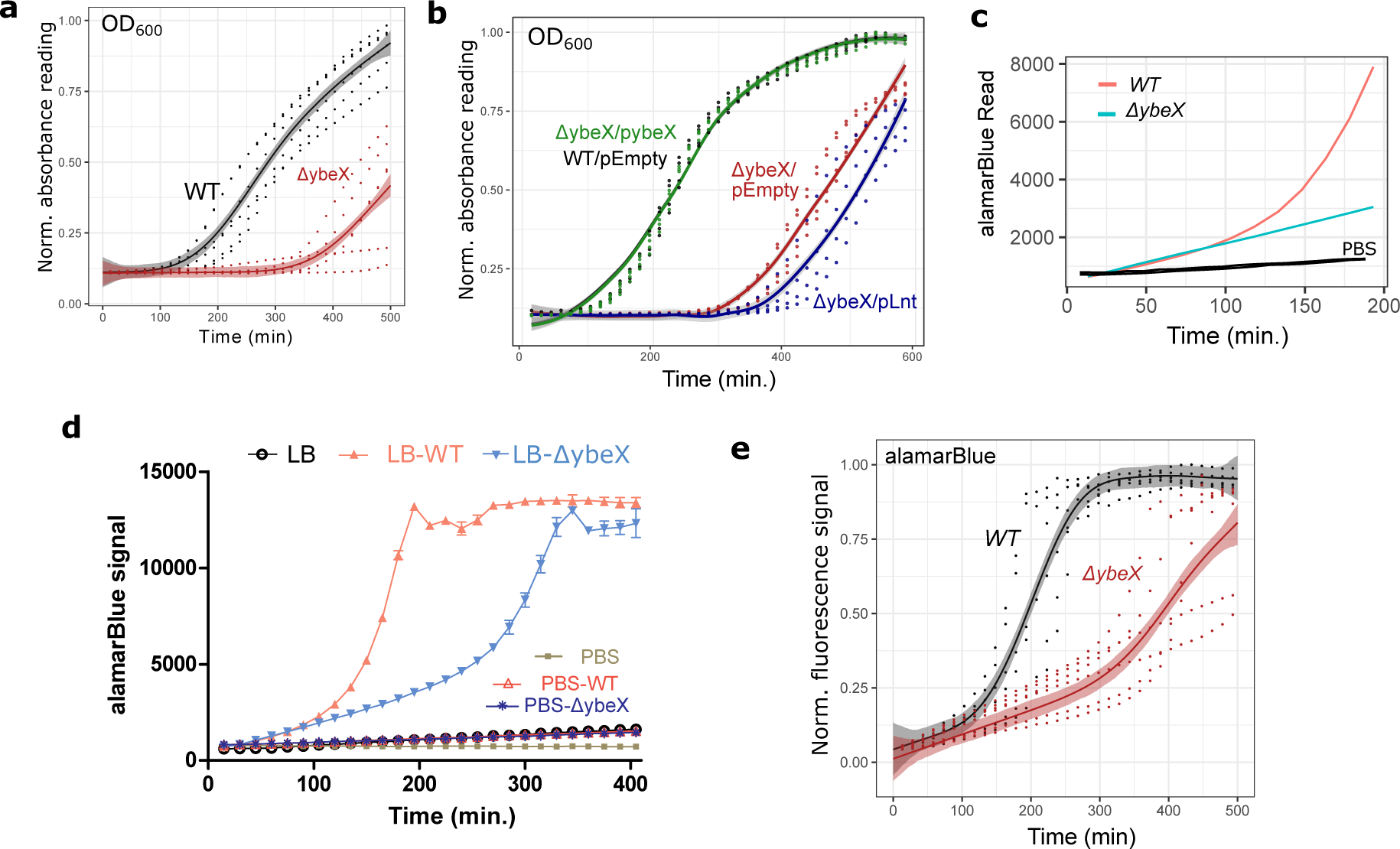
*ΔybeX* cells are metabolically active during the extended lag phase. (**a**) Overnight LB cultures of wild-type (WT) and *ΔybeX* strains were diluted, and regrowth was monitored in a 96-well plate reader. (**b**) Growth of *ΔybeX* strain overexpressing *ybeX* (pybeX) or *lnt* (pLnt) genes ectopically in comparison with wild type and *ΔybeX* strains harboring empty plasmid. (**c**) The Alamar Blue fluorescence signal of WT and *ΔybeX* strains was monitored in liquid LB medium and PBS buffer (the cells were diluted in 1xPBS, submerged black lines). (**d**) Fluorescence reading of Alamar Blue signal (excitation 545nm, emission 590nm) for sterile LB and PBS, or WT and *ΔybeX* cells diluted in LB (LB-WT and LB-*ΔybeX*) or sterile PBS (PBS-WT and PBS-*ΔybeX*). (**e**) The stationary phase outgrowth of wild type and *ΔybeX* strains in liquid LB medium where panel e shows Alamar Blue signal, indicating metabolic activity.

**Fig. S3.**
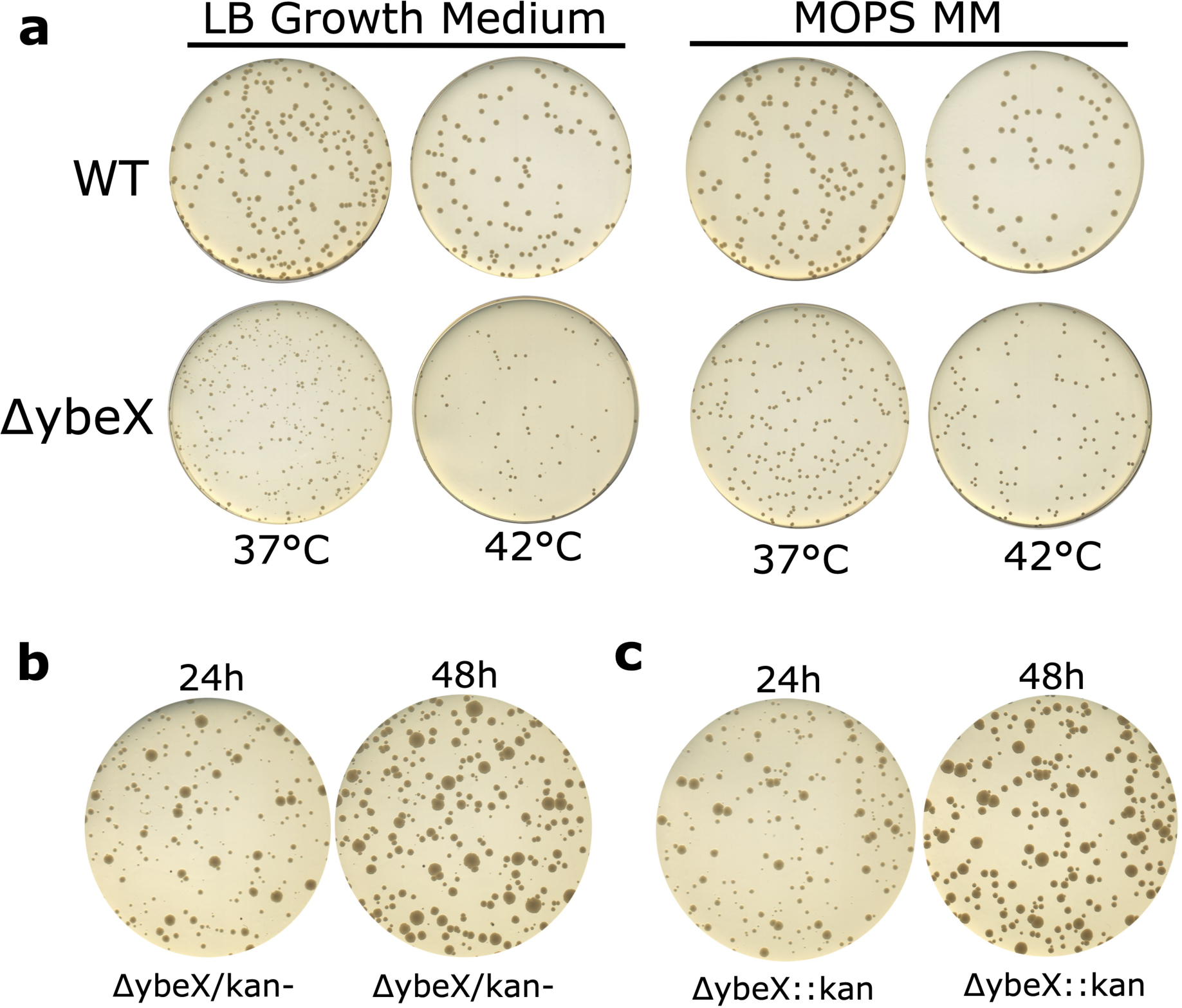
*ΔybeX* strain gives unequal colonies when plated directly from overnight cultures. (**a**) Colony inspection of WT and *ΔybeX* strains on LB agar plates precultured in LB and MOPS minimal media (MOPS MM, supplemented with 0.3% glucose as carbon source). A single colony for the indicated strains was aerated in a 125 mL glass flask in 10 mL of LB or MOPs for 16-18 hours at 37°C in a shaker incubator (200rpm). Overnight-grown saturated cultures were diluted and plated on LB Agar plates. The plates were incubated at indicated temperatures for 16 hours and then scanned. (**b**,**c**) A single colony of indicated strains were picked and inoculated into the LB liquid medium. The cells were grown at 37°C in a shaker incubator. The following morning the cells were diluted and plated on LB agar plates. The plates were incubated at 37°C. The same plate was scanned after 24 or 48 hours. The scanned plates are presented next to each other. The kanamycin resistance cassette is absent in *ΔybeX/kan-* strain.

**Fig. S4.**
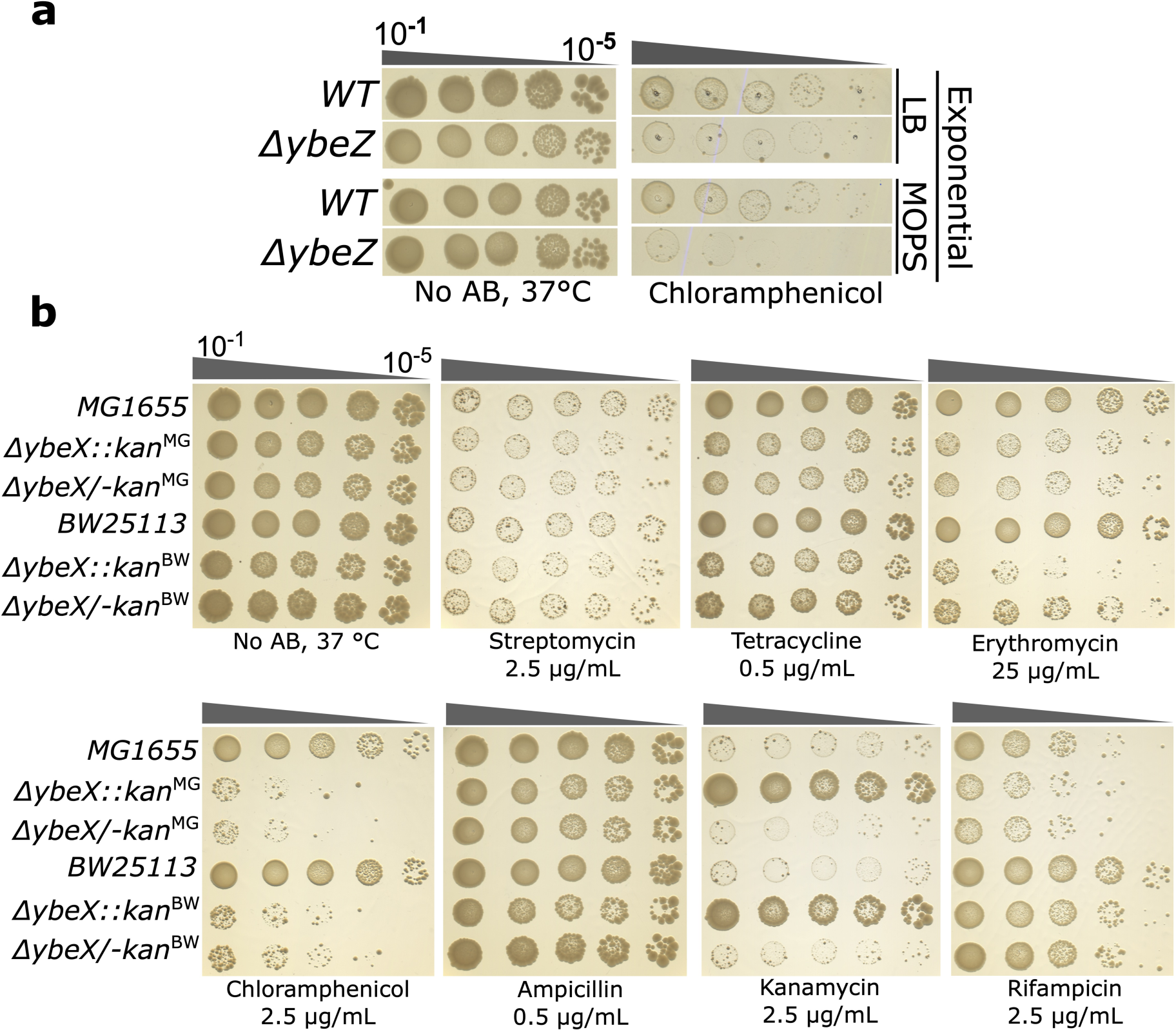
*ΔybeX* cells are sensitive to structurally unrelated ribosome-targeting antibiotics. (**a**) Dot-spot assay for Keio wild-type and *ybeZ* deletion strains on LB agar plates without any antibiotics (denoted as “No AB”) or 4 µg/ml chloramphenicol containing plates. The bacterial cells were grown in liquid LB or MOPS minimal medium overnight at 37°C. Next day, the cell cultures were diluted in fresh media and grown till OD_600_ reached 0.2-0.5. Exponentially growing cells were diluted and spotted. (**b**) Comparisons of two isogenic wild-type *E. coli* strains, one of which is MG1655 and the other BW25113, and the corresponding *ybeX* deletion strains *ΔybeX::kan^MG^* and *ΔybeX::kan^BW^*, respectively. Removal of kanamycin resistance cassette (*kan*) from *ΔybeX* in both *E. coli* strains, *ΔybeX/-kan^MG^* and *ΔybeX/-kan^BW^*, does not affect the cellular response to antibiotics.

**Fig. S5.**
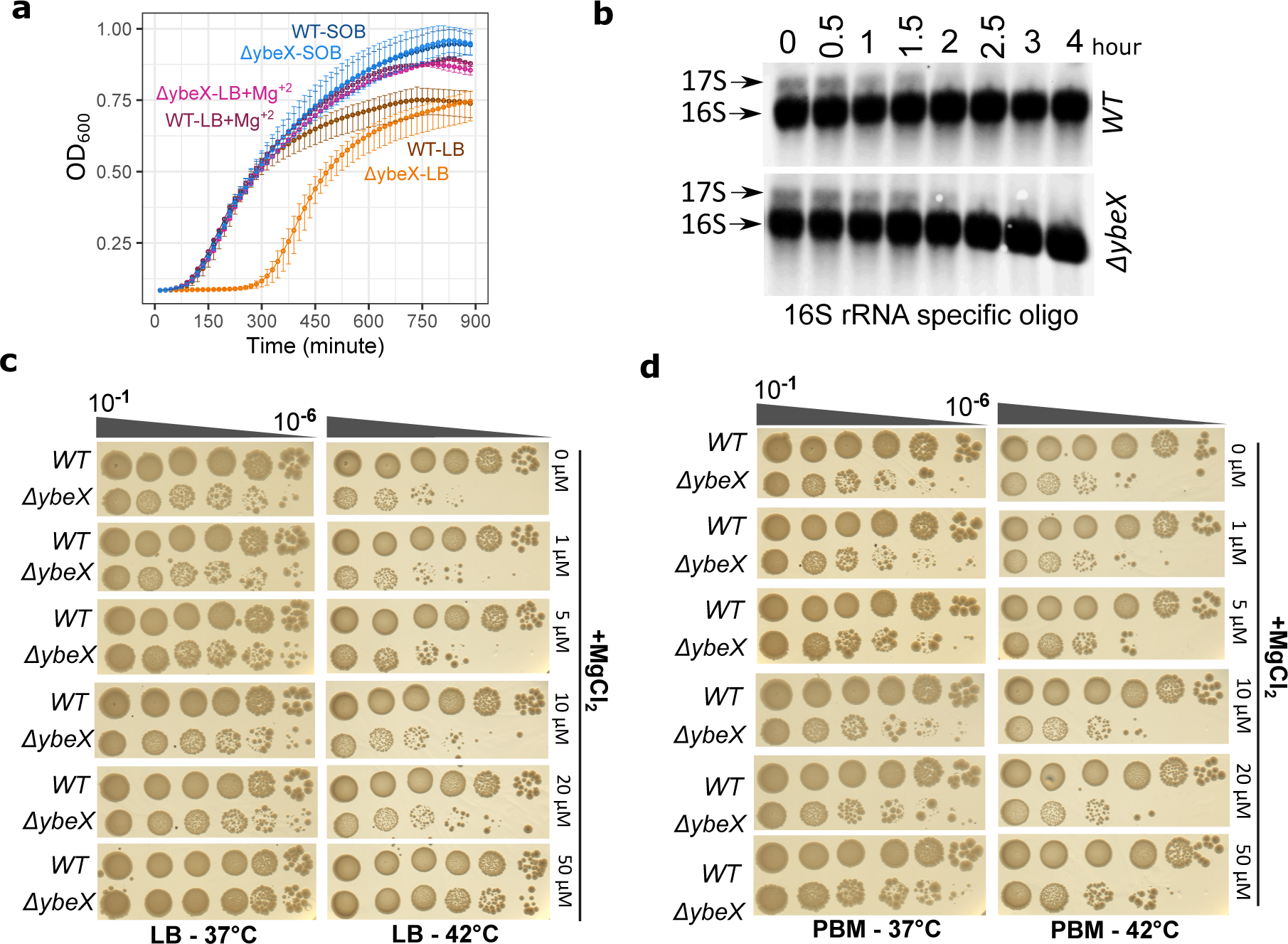
*ΔybeX* phenotypes vanish when the cells are grown in magnesium-rich complex growth media. (**a**) Bacterial growth was monitored by measuring optical density at 600 nm (OD_600_) for WT, and *ΔybeX E. coli* strains grown in LB, SOB or LB supplemented with 10 mM MgCl_2_. The cells were diluted in a 96-well plate containing LB liquid media. The means of three independent experiment are shown as dots, and the error bars are 95% CIs. (**b**) Northern Blot analysis of total RNA isolated from WT and *ΔybeX E. coli* strains grown in LB liquid media at 37°C under sufficient magnesium concentration. The first time point (0 hour) indicate the first sampling when the OD_600_ reached 0.3. The cells were pelleted and snap frozen at the indicated time points and stored at −80°C. The total RNA was purified using the hot-phenol extraction method. The Northern blotting was performed using 16S rRNA (position 338-356) specific Cy5 labelled DNA probe. (**c**-**d**) Wild-type and *ΔybeX* strains were grown overnight at 37°C with good aeration in LB (d) or PBM (e) liquid complex growth media with 0-50µM MgCl_2_ supplementation. Dilutions were spotted on LB agar plates and incubated at indicated temperatures, 37°C and 42°C.

